# Cross population comparison of complex migration strategies in a declining oceanic seabird

**DOI:** 10.1101/2023.06.01.541278

**Authors:** Nina J. O’Hanlon, Rob S.A. van Bemmelen, Katherine R.S. Snell, Greg J. Conway, Chris B. Thaxter, Helen Aiton, David Aiton, Dawn E. Balmer, Sveinn Are Hanssen, John R. Calladine, Sjúrður Hammer, Sarah J. Harris, Børge Moe, Hans Schekkerman, Ingrid Tulp, Elizabeth M. Humphreys

## Abstract

Human-driven change is impacting ecosystems globally, with consequential declines in biodiversity. Long-distance migrants are particularly susceptible as they depend on conditions over large geographical scales and are more likely to experience a greater range of pressures. One long-distance migrant that has experienced substantial declines across the North-East Atlantic is the Arctic Skua *Stercorarius parasiticus* however, little is known about their migratory behaviour or the routes they undertake. We used geolocators with salt-water switches to track Arctic Skuas from four breeding populations. To investigate migration strategies, we developed a Hidden Markov Model approach that used saltwater immersion data to classify stopovers and transit flights. We found that the skuas used several discrete staging areas during their south and northbound migrations with an area of apparent high marine productivity in the mid-North Atlantic being of high importance. Extensive overlap of individuals from different breeding populations occurred in staging areas, resulting in weak spatial connectivity between breeding and staging areas, further emphasizing their importance. At the population-level, variation in migration strategies was driven by individuals from Svalbard, which is declining less than the other populations tracked. Relative location of wintering areas also influenced migration strategies, with individuals migrating further spending a smaller proportion of their migration at stopovers compared to those wintering closer, and instead employed a fly-and-forage strategy. Identifying the complex non-breeding distribution, weak migratory connectivity and highly conserved migration strategies of Arctic Skuas is a vital step to link how conditions experienced during migration influences population dynamics and to prioritise future research and conservation actions.

## Introduction

Anthropogenic driven environmental change is causing large scale adverse impacts on ecosystems globally resulting in degraded habitats and changes in food availability (Kubelka et al. 2021). Long-distance migrants are particularly susceptible to encountering multiple stressors throughout the annual cycle and have experienced greater population declines than short-distance and resident species (Kubelka et al. 2021). Given that the different periods of the annual cycle are linked, environmental conditions that migrant species experience during the non-breeding season can influence their survival or condition, with carry-over effects on the demography and trends of breeding populations (Fretwell 1972, Marra et al. 1998, 2015, Harrison et al. 2011, Schultner et al. 2014, Fayet et al. 2017).

Having spatiotemporal predictable resources to exploit is important for long-distance migrants as the reliability of cues to track unpredictable, distant profitable conditions declines with greater distances from their breeding areas (Trierweiler et al. 2013, Brown et al. 2021). This can result in the migratory routes and strategies of long-distance migrants being more fixed than short-distance migrants, driven by knowledge learned from previous migrations or following conspecifics (Knudsen et al. 2011, Campioni et al. 2020, Brown et al. 2021). Areas of spatiotemporal predictable resources can therefore result in high concentrations of individuals from different populations and species in discrete locations (Davies et al. 2021).

Where different breeding populations of a species have shared wintering or staging areas spread over a large geographical area (weak migratory connectivity), populations may be buffered from adverse localised environmental conditions (Trierweiler et al. 2014, Knight et al. 2021). This may also result in similarities in demographic metrics, for example linked to shared foraging or weather conditions (Koenig & Liebhold 2016, Desprez et al. 2018). Alternatively, breeding populations can have spatially distinct wintering or staging areas (strong migratory connectivity; Palm et al. 2015) that can drive differences in breeding population trends, for example, if a high proportion of individuals from one population are negatively impacted by localised stressors that occur in their non-breeding location (Merkel et al. 2021). Given that many long-distance migrants concentrate in few, shared areas of high productivity during migration (Davies et al. 2021), it might be expected that populations of these species show weak connectivity to staging areas (Knight et al. 2021). Environmental change or human activities that reduces resource availability at staging areas can therefore be detrimental to several populations, particularly if alternative opportunities for safe refuelling along migration routes are limited (Knight et al. 2021).

In addition to which staging areas are used by specific breeding populations, understanding the migration strategy of long-distance migrants, i.e., the number of stopovers and the time spent at each area during a single migration (Bauer et al. 2016), is also important to determine the exposure of individuals to localised conditions, with consequences on population dynamics (Rakhimberdiev et al. 2018). Variation in migration strategies can be driven by where individuals breed and winter but also by the availability and quality of foraging habitats available along their migration routes (Newton 2008, Conklin et al. 2010) as higher energetic and fitness costs are associated with longer migrations (Pelletier et al. 2020, Buechley et al. 2021). Consequently, long-distance migrants can respond to higher energy requirements by either staging more frequently or for longer periods, or undertake a ‘fly-and-forage’ strategy, where individuals forage regularly whilst continuing to migrate over large distances (Strandberg & Alerstam 2007, Dias et al. 2012, Amélineau et al. 2021).

Many seabirds are long-distance migrants and are the most threatened group of birds globally (Dias et al. 2019). The Arctic Skua *Stercorarius parasiticus* has experienced long-term declines in breeding populations across the North-East Atlantic including in Scotland (Perkins et al. 2018), the Faroe Islands (Santos 2018), Norway (Henriksen & Hilmo 2015, van Bemmelen et al. 2021) and Iceland (Icelandic Institute of Natural History, 2018). These declines have resulted in the Arctic Skua being classified as Vulnerable on the European Red List of Species (BirdLife International 2021). In other areas of the North-East Atlantic the status of Arctic Skua is not clear, but the Svalbard population appears stable or weakly declining (Henriksen & Hilmo 2015). Declines have been associated with food shortages and predation during the breeding season influencing productivity as well as reduced survival of adults and immatures (Perkins et al. 2018, van Bemmelen et al. 2021, Santos et al. in prep.). However, given that Arctic Skuas spend a considerable amount of time away from the breeding area, conditions outside the breeding season are also likely to influence population trends, and there is evidence of time lag effects from wintering conditions on subsequent chick survival (Santos et al. in prep.). Arctic Skuas from breeding populations in the North-East Atlantic winter in multiple locations across the Atlantic Ocean and adjacent seas (van Bemmelen et al. 2023). In contrast, little is known about their migratory routes, staging areas or migratory strategies. Identifying these aspects of migration may help explain the differential population trajectories of Arctic Skuas in the North-East Atlantic and inform effective conservation actions (Marra et al. 2015, Dunn et al. 2019, Strøm et al. 2021).

The development and miniaturisation of modern biologging technology has enabled us to obtain more detailed spatial and temporal information on individuals across the annual cycle (Tuck et al. 1999, Wilson et al. 2002, Burger & Shaffer 2008, Strandberg et al. 2009). We used geolocators to compare migration strategies and routes used among populations during the post-breeding southbound (autumn) migration and the pre-breeding northbound (spring) migration. Using saltwater immersion data, we developed a method to categorise when migrating Arctic Skuas were on transit flights or stopovers, defined as locations where individuals typically stayed for short durations to rest or feed (Warnock 2010). These stopover locations were used to identify the core staging areas of individuals and to test the connectivity of breeding populations to staging areas. Due to the importance of discrete areas of high marine productivity to migrating seabirds (Davies et al. 2021), and that Arctic Skuas show weak migratory connectivity between breeding and wintering areas (van Bemmelen et al. 2023), we predict that the mixing of individuals from different breeding populations occurs before individuals arrive at their wintering areas, and so that, from a breeding population perspective, Arctic Skuas show weak migratory connectivity to staging areas. Given the weak migratory connectivity between breeding and wintering areas, we also expect that migration routes and strategies of individuals from the same breeding population will differ. Specifically, we predict that individuals migrating between more disparate breeding and wintering areas will make longer transit flights and therefore require more and/or longer stopovers than individuals wintering closer to the breeding colony.

## Methods

### Study species / region

Adult Arctic Skuas were caught during the breeding season between 2009 and 2019 from four breeding populations in the North-East Atlantic: Scotland, Faroe Islands, mainland Norway and Svalbard (Table 1, S1, Figure 1). Individuals were trapped on the nest using walk in traps and remote release nooses or away from the nest using net guns. A leg-mounted geolocator, attached to a plain or unique alpha-numeric plastic ring, was deployed on each individual. The combined weight of the ring, geolocator and attachments was *c*. 2.5g. The body mass at deployment of individuals from which geolocators were retrieved was 447.0 ± 48.8g; N = 108. Therefore, deploying the ring and geolocator added less than 1% of the skuas’ body weight. In addition, eight Scotland skuas (four from Fair Isle and four from Rousay) also had glue-mounted GPS devices temporary attached to their back feathers for up to three weeks post-capture (maximum combined geolocator, GPS and attachment weight of 12.9g, range 2.33 – 3.27% of body mass). All GPS devices fell off before their southbound migration and we do not believe they affected the migratory behaviour of these individuals. Tagged individuals were recaptured at the breeding colony between 2010 and 2021 to retrieve geolocators.

**Table 1.** Number of years individual Arctic Skuas were tracked for, by population.

|  | Faroe<br>Islands | Norway | Scotland | Svalbard | Total |
| --- | --- | --- | --- | --- | --- |
| Number of years individuals<br>were tracked for |  |  |  |  |  |
| 1 | 19 | 27 | 6 | 17 | 69 |
| 2 | 4 | 17 | 4 | 5 | 30 |
| 3 | 0 | 11 | 0 | 10 | 21 |
| 4 | 0 | 2 | 0 | 5 | 7 |
| 5 | 0 | 0 | 0 | 3 | 3 |
| 6 | 0 | 0 | 0 | 1 | 1 |
| Total number of individuals | 23 | 56 | 10 | 40 | 129 |
| Total number of tracks | 27 | 102 | 14 | 98 | 241 |
| Total number of tracks with<br>saltwater immersion data | 27 | 98 | 10 | 74 | 209 |

**Figure 1.**
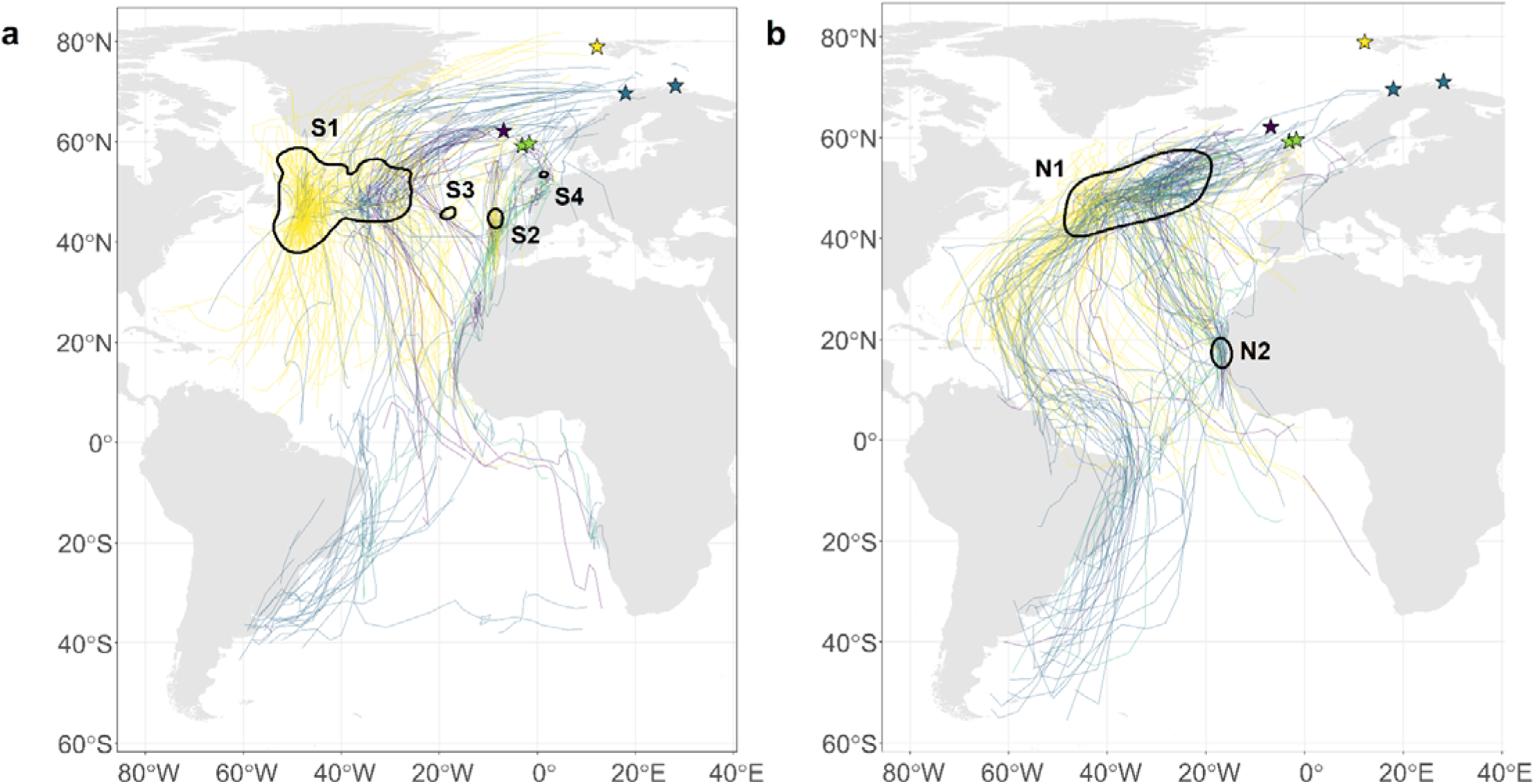
Core staging 50% utilisation distribution (UD) kernels (black polygons) for tracked Arctic Skuas during a) southbound and b) northbound migration, combining all locations classified as stopovers (from the 2-state HMM using saltwater immersion data, see main text for details), from Scotland (green, 10 tracks); Faroe Islands (purple, 27 tracks); Norway (blue, 98 tracks); and Svalbard (yellow, 74 tracks). Staging areas are labelled S1, S2, S3 and S4 for southbound migration and N1 and N2 for northbound migration. Tracks show smoothed migration routes of Arctic Skuas from each population to illustrate the broad-scale connectivity between breeding, core staging and wintering areas (see figures S1a-d for smoothed migration routes by population and wintering area). The UD kernel labels match those in Table S7. Stars depict breeding colonies: Scotland (green); Faroe Islands (purple); Norway (blue); and Svalbard (yellow); with individuals tracked from two colonies in Scotland and Norway.

Three types of geolocators were deployed, produced by British Antarctic Survey (mk9, mk13. mk15, mk18h), Biotrack (mk3006, mk4083) and Migrate Technology (C65, C65s, C250) (Table S2). The majority of geolocators (156, 87%, of the 177 deployed) also recorded conductivity, or saltwater immersion data. These recorded when the device was submerged, which can be used as a proxy of behaviour during migration (Lecomte et al. 2010). The different geolocator models recorded saltwater immersion data differently and sampled: bursts of measurements every 3 seconds (sec) with number of sample bursts recorded every 5 or 10 minutes (min); or bursts of 6 sec and recorded on change of state from wet to dry or dry to wet (Table S3).

### Data processing

Data downloaded from retrieved geolocators were processed using R 4.0.3 (R Core Development Team 2020, Figure S1). Initial processing to calculate sunrise and sunset times (twilight events) was conducted using the *twilightCalc* function in the *GeoLight* R package (Lisovski & Hahn 2012) using a light-level threshold of 2 (for the Migrate Technology geolocators) or 10 (for the British Antarctic Survey and Biotrack geolocators). Based on twilight events and sun elevation angles, longitudes were calculated from the timing of local noon and midnight, and latitudes from day length. The mean error of geographical positions estimated from geolocators is ±185 km (Phillips et al. 2004). This error increases for latitude during the equinoxes when daylength is similar across the world (Frederiksen et al. 2012), therefore we removed positions within 17 days of the north and southbound equinoxes (20 March and 22 September).

Geographic positions outside the breeding area were assigned as either on southbound migration, at the wintering area or on northbound migration. Departure and arrival from and to the breeding and wintering areas were identified from the raw position estimates, in particular longitude, which can still be reliably estimated within the equinox periods. Onset of migration was typically shown as a rapid movement (often with a strong easterly or westerly direction) away from the breeding or wintering area. The area skuas were located between the end of the southbound migration but before the onset of the northbound migration was defined as the wintering area (van Bemmelen et al. 2023). Six main wintering areas were identified for Arctic Skuas breeding in the North-East Atlantic based on January mean positions: Mediterranean Sea, Canary Current, Caribbean region, Gulf of Guinea, Benguela region and Patagonian Shelf (van Bemmelen et al. 2023). The period from when an individual departed the breeding area and arrived at the wintering area was defined as the southbound migration, whilst the period between departing the wintering area and returning to the breeding area was defined as the northbound migration. Positions from the breeding season and wintering areas were excluded from the analyses. Thirty-eight geolocators failed or ran out of battery before individuals returned to breeding areas, therefore, these tracks were only included in the relevant analyses where data was available for a complete southbound migration.

### Activity derived from saltwater immersion data

All saltwater immersion data from the different geolocator models was derived to the same sampling frequency of 3 sec resulting in summed values between 0 and 200 at the end of each 5- or 10-min period. From this data, the activity patterns of individuals during migration were identified (see Lecomte et al., 2010). When the device was dry (value of 0) we assumed an individual was on a transit flight, whereas when the device was wet (value of 200) we assumed an individual was resting on the water. For intermediate values (1 – 199) we assumed the skuas were foraging, however given that Arctic Skuas frequently kleptoparasitise, dry periods may also include some wholly aerial foraging activity, although this will likely only make up a small proportion of this time.

To determine the activity of skuas during migration we estimated the proportion of 5- or 10- min periods in each day (24-hour period including daylight and darkness) that were categorised as dry (as a proxy for active migration). As the saltwater immersion data recorded by the geolocators was not influenced by the equinoxes this provided data for the entire south and northbound migration. We therefore used the saltwater emersion data to classify transit flight and stopover locations independent of geographical positions.

To classify transit flight or stopovers we carried out Hidden Markov models (HMMs, McClintock & Michelot 2021). We fitted a two-state HMM with a single data stream, the proportion of each day that was dry, using the R package *momentuHMM* (McClintock & Michelot 2021). We assumed a two-state model based on the knowledge that transit flights would be associated with days where a greater proportion of the day was recorded as dry, and that stopovers would be associated with days where a greater proportion of the day was recorded as wet or intermediate (Figure S2). A beta distribution was used for the dryness data as it is proportional data that is truncated between 0 and 1 (with 1 reflecting an entirely dry day or 24-hour period). For tracks with saltwater immersion data, this resulted in all migration days being assigned to one of two states. The first state reflected a higher proportion of the day recorded as dry (mean 0.50 ± SD 0.20), which we interpreted as transit flights, and the second state reflected a lower proportion of the day recorded as dry (mean 0.13 ± SD 0.09), which we interpreted as stopovers. Therefore, a stopover could be as short as a single day. To check the classification of positions, we compared travel rates (km/day) during transit flights and stopovers during south and northbound migrations. We calculated travel rates per day by measuring the daily distance travelled between double smoothed positions (see below), using the *disthaversine* function in the *Geosphere* R package (Hijmans 2019). We ran a general linear model with travel rate (km) as the response variable and state (transit flight or stopover) as a fixed effect. Travel rate was natural logarithm transformed to meet the assumptions of normality. Travel rates were significantly greater during transit flight (408 ± 237 km) than at stopovers (102 ± 50 km).

### Analysis of geolocator data

#### Visualisation of migration routes

To determine the routes of Arctic Skuas during their south and northbound migrations we used ‘double smoothed’ positions to smooth some of the error around the raw positional data (Gilg et al. 2013). This involved firstly, taking the average latitude and longitude of the two daily positions and secondly, calculating 3-day running means from these daily averaged positions using the equations used by Gilg et al. (2013) for Long-tailed Skuas *Stercorarius longicaudus*. As positions 17 days either side of the equinoxes were removed, sections of the track either side of each equinox were smoothed separately, resulting in gaps in the routes when the equinox occurred during migration (see Table S4 for the proportion of tracks and migration duration that overlapped with the equinoxes). Smoothed routes provide a broad pattern of migration movement rather than fine-scale routes taken by individuals. Given the large number of tracks, south and northbound migration routes are displayed in Figure S3 by breeding population and wintering area.

#### Identification of core staging areas during migration

To identify core staging areas we included all tracks with available activity data to create utilisation distributions (UD) kernels, using ‘double-smoothed’ positions classified as stopovers by the HMM outside the equinoxes when positional data was available. Kernels were created for both the south and northbound migration, weighted by the number of tracks per population, using a Lambert azimuthal equal-area projection and 50 km grid cells. As we were interested in staging areas at the species-level, we created UD kernels across populations and years. There was a high degree of overlap between the species-level UD kernels and those created for each population and year separately (see supplementary material, Figures S4, S5, S6, Tables S4, S5). We calculated the most appropriate smoothing parameter (*h*) for each population and season using a custom function in R that derives a “minimum” (or adjusted) *h*-reference bandwidth to avoid potential issues of over or under-smoothing. This method searches iteratively for the smallest *h* over progressively smaller scales starting with the *h*-reference bandwidth value and selects the smallest *h* prior to the eventual break-up of the 95% KDE spatial polygons. Therefore, different *h* values were used for each population and migration period (southbound: Scotland = 445652; Faroe Islands = 336202; Norway = 420197; Svalbard = 131169. Northbound: Scotland = 356356; Faroe Islands = 363932; Norway = 213437; Svalbard = 232252). As we were interested in the core staging areas used by the skuas, we calculated the 50% UD kernel contour for each migration period using the *kernelUD* function in the *adehabitatHR* R package (Calenge 2006, 2015). To identify the proportion of individuals that used each species-level core staging area during each migration (southbound, northbound) we extracted the stopover positions that contributed to each area using the *over* function in the *rgeos* R package (Bivand & Rundel 2019). To highlight the importance of each staging area we also calculated the proportion of stopover days (bird days) individuals spent within each staging area by breeding population and wintering area (Figure 2).

**Figure 2.**
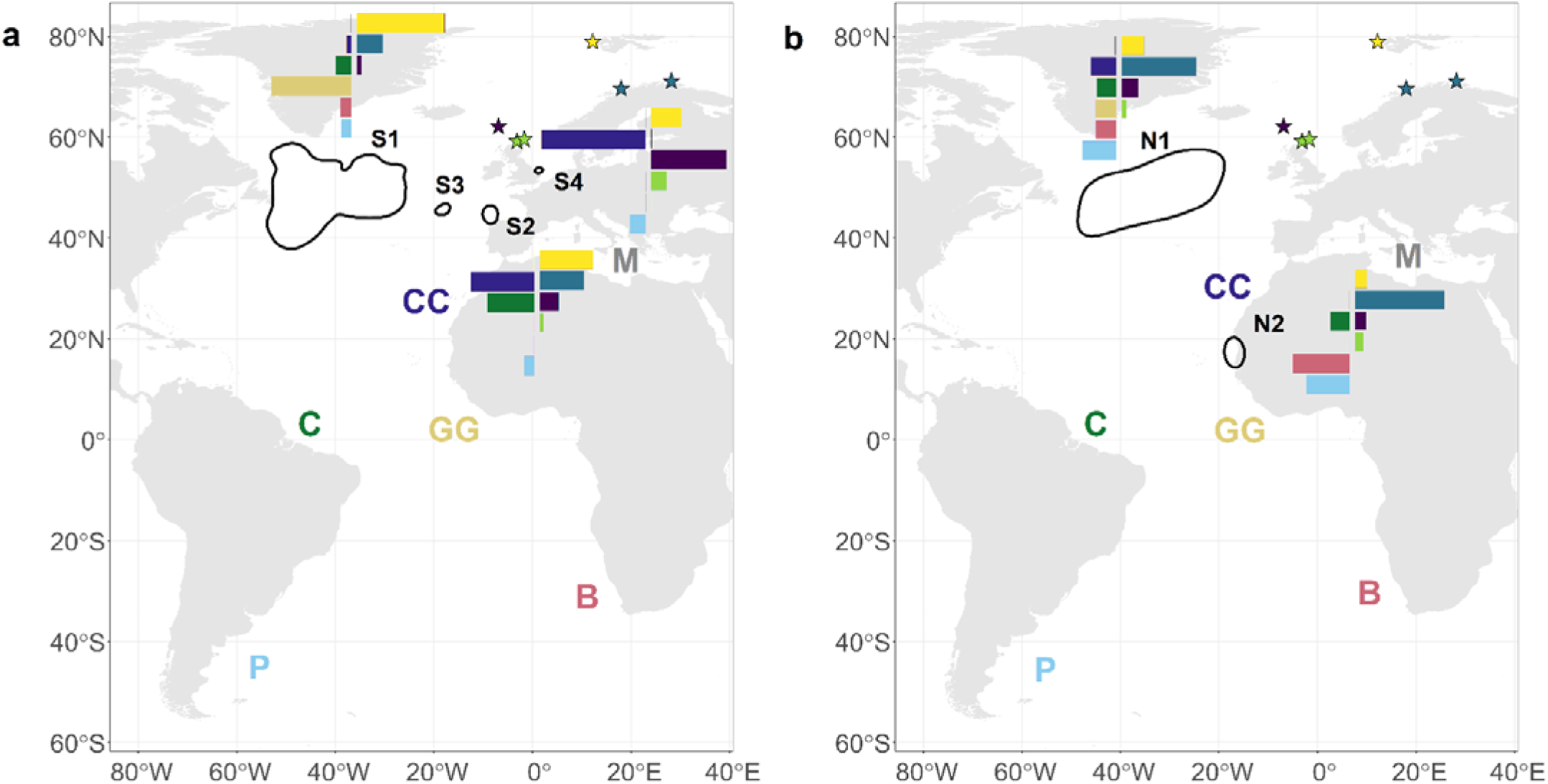
Core staging 50% utilisation distribution (UD) kernels (black polygons) for tracked Arctic Skuas during a) southbound and b) northbound migration, combining all locations classified as stopovers, from Scotland (green stars); Faroe Islands (purple star); Norway (blue stars); and Svalbard (yellow star). For each staging area, the right-hand bars show the proportion of bird days by breeding population that staged within that core 50% UD. The left-hand bars show the proportion of bird days spent within each core 50% UD by wintering area. The bars are on a scale from zero to one, with the total proportion for each breeding and wintering area across staging areas adding up to one. The number of bird days associated with each staging area during southbound migration was: S1 = 1562, S2 = 38, S3 = 15, S4 = 8; and for northbound migration: N1 = 1124 and N2 = 102. Bars are not shown for southbound staging area S3 as this only involved two individuals from Svalbard that wintered in the Caribbean region. The UD kernel labels match those in Table S7. The coloured letters refer to the main wintering areas: M = Mediterranean Sea, CC = Canary Current, C = Caribbean region, GG = Gulf of Guinea, B = Benguela region and P= Patagonian Shelf. See van Bemmelen et al. (2023) for details on wintering areas.

To determine the overlapping space use of the 50% UD kernels between south and northbound migrations we calculated Bhattacharyya’s affinity index (BA, Bhattacharyya 1943; Fieberg and Kochanny 2005) using the *kerneloverlaphr* function in *adehabitatHR* (Calenge 2015). The BA index provides a measure of similarity between two UD kernels between 0 and 1, with 0 indicating no overlap and 1 being identical (Fieberg & Kochanny 2005).

#### Migratory connectivity of breeding populations to staging areas

To quantify the migratory connectivity of Arctic Skuas between breeding populations and staging areas we calculated Mantel correlation coefficients (r_M_) for both south and northbound migrations, using the *estMantel* function in the R package *MigConnectivity* (Cohen et al. 2017). Mantel correlation coefficients (r_M_) range from -1 to 1, with values around zero or lower indicating low migratory connectivity, and values towards 1 indicating high migratory connectivity (Ambrosini et al. 2009, Cohen et al. 2017). As some individuals stopped at more than one staging area during their south and northbound migrations (see results), we selected the longest staging period of each migration. Across breeding populations, 46 individuals had data on migratory routes for multiple years. Therefore, for these individuals we only included the first south and northbound migration in this analysis. Based on dynamic time warping (DTW) ratios (see Appendix), migration routes were significantly more similar within-individuals than between-individuals, during both south and northbound migrations, indicating high repeatability of routes by individuals between years.

#### Population specific variation in migration strategies

To test whether skuas that migrated to more distant wintering areas made more and longer stopovers and transit flights (as classified by the two-state HMM), for each track (where saltwater immersion data was available) and migration (southbound and northbound) we calculated 1) the number of stopovers and transit flights, 2) the duration in days of each stopover and transit flight, and 3) the total number of days classified as stopover or transit flight per migration (summarised for each population: Table S8; and for each wintering area: Table S9).

### Statistical analysis

To test for variation among breeding populations and wintering areas in the migration strategies of Arctic Skuas, we ran two separate generalised linear mixed models (GLMMs) with 1) the total duration of stopovers and 2) the proportion of each migration spent at stopovers as the response variables. We focused on total duration (in days) as this value captured, and was significantly correlated with, both the number and mean duration of stopovers (Number: t = 10.98, p < 0.001, r = 0.50; Mean duration: t = 16.39, p <0.001, r = 0.65; N = 370).

Interactions between population and migration period, and between wintering area and migration period, were included as fixed effects with individual ID as a random effect to account for where we had multiple tracks per individual. Models were run in the *glmmTMB* R package (Brooks et al. 2017) with a Gamma distribution and log link function for the total duration (in days) of stopovers, and a beta distribution and a logit link function for the proportion of migration at stopovers.

Tukey adjusted post-hoc multiple comparisons were carried out in the R package *emmeans* (Lenth et al. 2021). All statistical analyses were performed in R, Version 4.0.3 (R Core Development Team 2021), with figures created using *ggplot2* (Wickham 2016). Throughout we report means and standard deviations (SD).

## Results

Across the four Arctic Skua breeding populations we retrieved data for 241 tracks, 209 of which also had saltwater immersion data, from 129 individuals, with 62 individuals tracked over multiple years (2 – 6 years; Table 1). As we focused on the Atlantic Ocean, one track from Norway of a bird over-wintering in the Indian Ocean was excluded from the analysis (van Bemmelen et al. 2023).

### Migration routes

The smoothed migration routes revealed that Arctic Skuas from the Faroe Islands, Norway and Svalbard typically headed west/southwest after the breeding season into the mid-Atlantic before travelling further south (Figure 1, Figures S3a-d). Conversely, individuals from Scotland largely departed to the south, with a small proportion of all tracks transiting through the North Sea and English Channel (southbound: minimum proportion of tracks = 0.07, N = 21; northbound: minimum proportion of tracks = 0.02, N = 4). During northbound migration, most individuals from all populations (minimum proportion of tracks = 0.90, N = 147) returned to their breeding colonies via the mid-Atlantic.

### Core staging areas during migration

Four core staging areas were identified during southbound migration, and two during northbound migration based on the stopover positions across all populations and years (Figure 1). During southbound migration over half (58%) of individuals used one of the identified staging areas, with 3% using two staging areas (Table S7). During northbound migration, 76% of individuals used one staging area, with 11% associated with two staging areas. The largest staging area, used by the skuas during both south and northbound migration, was located along the mid-Atlantic ridge (Kernels S1 and N1, Figure 1), which accounted for the high spatial overlap between the south and northbound staging areas (Bhattacharyya’s affinity = 0.76). During southbound migration, at least 50% of Arctic Skuas stopped over within the mid-Atlantic staging area, all of which were from the three more northern populations (Faroe Island, Norway and Svalbard; Figure 2, Table S7). Three smaller staging areas, used by a small number of individuals from all populations, were identified off the east coast of England, off northwest Spain and in the Atlantic between the northwest Spain and mid-Atlantic staging areas (Figure 2, Table S7). During northbound migration at least 83% of individuals had stopovers within the mid-Atlantic staging area, from all four populations (Figure 2, Table S7). A smaller northbound staging area was also identified off West Africa, used by at least 14% of individuals (N2, Figure 2).

### Migratory connectivity of breeding populations to staging areas

The high degree of mixing of individuals within the core staging areas was reflected by low Mantel correlation coefficients (r_M_) for both the south (r_M_ = 0.25, 95% Confidence Interval: 0.06 - 0.46; n = 72) and northbound (r_M_ = 0.20, 95% Confidence Interval: 0.02 - 0.37; n = 97) migration, indicating low spatial migratory connectivity. Although migratory connectivity between breeding and species-level staging areas were low across populations, individuals from Svalbard, the most northerly breeding population, tended to stage further west in the North Atlantic than individuals from elsewhere, especially during southbound migration (Figure 1, S4).

### Population specific variation in migration strategies

There was evidence for some variation in migration strategies among spatially discrete breeding populations and wintering areas (Figure 3). For the total duration of days spent at stopovers, there was a significant interaction between breeding population and migration period (χ^2^ = 15.76, p = 0.001; Figure 3a), with substantially fewer days at stopovers during northbound than southbound migration among individuals from Svalbard (t_348_ = -5.64, p < 0.001), but not in the other populations (p > 0.36). During northbound migration, individuals from Svalbard also spent significantly fewer days at stopovers compared to those from the other populations (p < 0.01; Figure 3a). The interaction between population and migration period for the proportion of migration spent at stopovers was also significant (χ^2^_5_ = 13.24, p = 0.0014; Figure 3b), with individuals from Svalbard spending a significantly lower proportion of time at stopovers during northbound than southbound migration (t_348_ = -5.64, p < 0.003), but again not in the other populations (p > 0.19). During northbound migration, individuals from Svalbard also spent a lower proportion of days at stopovers compared to individuals from Norway and the Faroe Islands (p < 0.001; Figure 3b).

**Figure 3.**
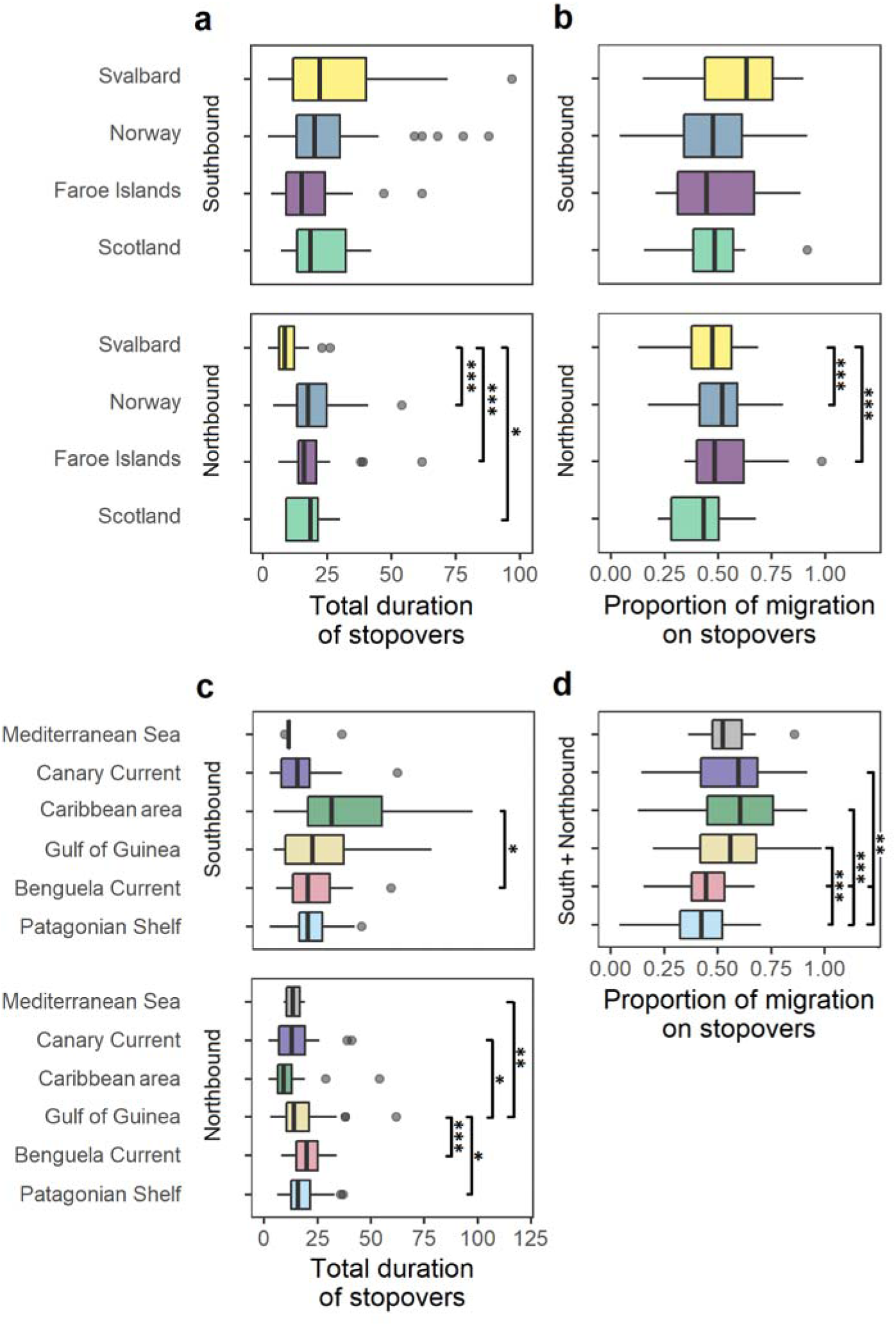
Boxplots showing the variation in staging migration strategies of tracked Arctic Skuas during both south and northbound migration, based on a) and c) the total duration (days) of stopovers, and b) and d) the proportion of days spent at stopovers, as a function of a) and b) breeding population or c) and d) wintering area. Breeding populations and wintering areas are ordered by highest to lowest latitude. Lines link populations that were significantly different to one another based on the results of Tukey adjusted post-hoc multiple comparisons of generalised linear mixed models (see main text for details). There was no significant interaction between wintering areas and migration period (southbound, northbound) for the proportion of days spent at stopovers. Asterisks refer to significance levels: ≤ 0.05*, ≤ 0.01**, ≤ 0.001***. Boxplots show median (horizontal line), inter-quartile ranges (box), and minimum and maximum values (whiskers).

Based on where individuals wintered, there was a significant interaction between wintering area and migration period for the total duration of days spent at stopovers (χ^2^_5_ = 20.39, p = 0.001; Figure 3c), with individuals wintering in the Caribbean region and off the Patagonian Shelf spending fewer days at stopovers during northbound compared to the southbound migration (p < 0.01), but not in the other populations (p > 0.36). During northbound migration, the number of days spent at stopovers by individuals migrating to the Gulf of Guinea also differed, with individuals spending more days at stopovers than those wintering in areas closer to their breeding population but fewer days than those at the furthest wintering areas (the Benguela Current and Patagonia Shelf; Figure 3c). There was also a significant influence of wintering area on the proportion of migration spent at stopovers (χ^2^_5_ = 46.90, p < 0.001; Figure 3d) but not interacting with migration period (χ^2^_5_ = 4.49, p = 0.48), with individuals migrating to the Benguela Current and Patagonia Shelf spending a lower proportion of days on migration at stopovers compared to individuals wintering closer to breeding areas.

## Discussion

Tracking Arctic Skuas with geolocators revealed that individuals from multiple breeding populations across the North-East Atlantic migrated across a vast area of the Atlantic Ocean, spanning both hemispheres. Through combining positional data with saltwater immersion data, we identified several discrete staging areas, and the migration strategies, used by individuals during migration.

### Migration routes

Arctic Skuas showed variation in migration routes among breeding populations driven by where the skuas staged, in addition to where they bred and wintered. Following the breeding season, individuals from the Svalbard, Norway and Faroe Islands breeding populations migrated westwards to stage along the mid-Atlantic ridge before continuing further south. Conversely, individuals from the most southerly population, Scotland, largely took a more easterly route around Britain, migrating south through the North Sea and English Channel along the Iberian Peninsula and West Africa. Based on a concurrent study which used Global Positioning System tags, the Scottish population are known to forage in the North Sea during the breeding season, (Johnston et al. in prep), which may have influenced their movements post-breeding.

During northbound migration, the majority of individuals from all populations headed to the mid-Atlantic ridge before returning to their respective breeding colonies. After leaving wintering areas in the southern Atlantic, a high proportion of individuals migrated north via West Africa, with birds that wintered in Patagonia diverging on their northbound migration between the Caribbean and West Africa. These south and northbound migration routes are similar to those of Arctic breeding Long-tailed Skuas, and of Arctic Tern *Sterna paradisaea* and Sabine’s Gulls *Xema sabini* that the skuas likely kleptoparasitise on route (Egevang et al. 2010, Stenhouse et al. 2012, Gilg et al. 2013). Following the routes of other seabirds such as small gulls and terns likely provide foraging opportunities inside and outside core staging areas (Paterson 1986, Furness 1987, Wuorinen 1992, Belisle & Giroux 1995, Gilg et al. 2013).

This is the first time the migratory routes of Arctic Skua in the North-East Atlantic have been identified, which is important given they spend a considerable proportion (70% of days on average across tracked individuals) of the annual cycle away from their breeding areas, where most current research effort has been focused (Perkins et al. 2018, van Bemmelen et al. 2021). This improved knowledge of where Arctic Skuas are distributed during migration can facilitate future efforts to identify stressors individuals may encounter along migration corridors, such as interactions with fisheries, including bycatch, and renewable energy developments. A potential threat to individuals migrating through the North Sea is posed by offshore wind farms, with Arctic Skuas being at relatively high risk to collision with turbines (Furness et al. 2013; Chirosca et al. 2022). This largely involved individuals from Scotland and the Faroe Islands during their southbound and, to a lesser extent, northbound migrations. Although, individuals from other populations, especially those wintering around the Canary Current and Mediterranean Sea that migrate through the North Sea instead of heading to the mid-Atlantic ridge, may also be at risk.

### Core staging areas during migration

The majority of tracked Arctic Skuas staged in an area east of the Grand Banks of Newfoundland to the mid-Atlantic Ridge, an area of tremendous conservation value that is used by many seabird species during south and northbound migration due to its temporally stable, high biological productivity (Davies et al. 2021). However, this is the first time this area has been shown to be used by migrating Arctic Skuas. Given the abundance of other seabirds, this area likely provides abundant foraging opportunities for the skuas and is likely critical during northbound migration to provide individuals with resources to optimise body condition before the breeding season (Belopolskii 1961). The importance of the mid-Atlantic ridge has led to part of this region being designated as a Marine Protected Area, which should help mitigate threats facing species that use it (North Atlantic Current and Evlanov Sea-basin MPA; Davies et al. 2021). However, this protected area only covers 20% and 22% of the mid-Atlantic staging area used by the skuas during south and northbound migration respectively. Individuals also staged at a smaller area associated with upwellings and high marine productivity off West Africa and around the Canary Current during their northbound migration, frequently used by other migrant and resident seabird species (Camphuysen & Van Der Meer 2005, Stenhouse et al. 2012, Grecian et al. 2016, Harrison et al. 2021). This staging area is also likely important in providing opportunities for the skuas to obtain resources during migration, and given the importance of this area for multiple seabird species, could also be considered for MPA designation (Lascelles et al. 2016).

### Migratory connectivity of breeding populations to staging areas

Arctic Skuas breeding in the North-East Atlantic overlapped extensively during their south and northbound migration at staging areas, showing weak spatial migratory connectivity similar to that between the breeding and wintering areas (van Bemmelen et al. 2023). The strength of migratory connectivity therefore declined once individuals departed from their breeding areas, much earlier than when they arrived at their wintering areas. This emphasises the importance of the core staging areas for individuals and populations from multiple breeding populations and the importance of quantifying migratory connectivity during migration in addition to wintering areas (Knight et al. 2021).

Understanding migratory connectivity throughout the annual cycle can help identify drivers of variation in population trends and prioritise where in time and space to focus conservation efforts (Palm et al. 2015, Rushing et al. 2016, Knight et al. 2021). The observed low migratory connectivity, particularly in association with populations being spread over a large geographical area, may suggest that Arctic Skuas will be buffered against adverse conditions or threats in the non-breeding season acting at local or regional scales, as well as being more resilient to environmental change (Trierweiler et al. 2014, Gilroy et al. 2016).

Conversely, changes in environmental conditions in core staging areas, for example driven by over-fishing and climate change, may lead to reduced food availability that has the potential to adversely impact multiple Arctic Skua breeding populations around the North Atlantic, as well as that of many other seabirds. Furthermore, extreme weather events in such key non-breeding locations may impact the survival rates and therefore trajectories of multiple populations (Reiertsen et al. 2021). It is also important to establish whether climate change leads to other marine areas becoming more productive, resulting in staging areas becoming more transient, to determine whether dynamic marine protected areas should be considered in future (Cashion et al. 2020), with potential consequences on future migration strategies and routes (Fayet et al. 2017, Lamb et al. 2017).

Here we focused on spatial migratory connectivity to understand the importance of the shared staging areas to the different breeding population. However, the temporal overlap of individuals from multiple populations at these locations can also be important given that conditions and threats often vary in time as well as space (Cohen et al. 2018, Knight et al. 2021). This may be particularly important for understanding how the mixing of high concentrations of seabirds, especially long-distance migrants from multiple populations, may facilitate the spread of disease, such as highly pathogenic avian influenza (Caliendo et al. 2022). Despite missing data around the equinoxes, our data suggests that although the skuas used similar, discrete staging areas during migration there was some temporal segregation, especially by individuals from Svalbard (Figure S7), attributed to their later migration schedules compared to the other populations (van Bemmelen et al.2023). Svalbard individuals visited staging areas considerably later than skuas from other breeding areas during south and northbound migrations. During southbound migration, Svalbard individuals also used a more westerly region of the mid-Atlantic staging area than other populations, which may be related to their later migration timings, and potential differences in where prey was concentrated at that time.

### Behavioural classification of migratory activity

By using a two-state HMM we successfully developed an approach to use saltwater immersion data to classify the activity of the Arctic Skuas during migration, specifically when individuals were in flight versus resting or foraging, even during the equinoxes where we had gaps in geographical locations. Although the skuas spent significantly less time in flight whilst at stopover locations than on transit flights, the proportion of time in flight was low for both activities. Therefore, even during transit flights the skuas spent part of the day resting or foraging. Part of this time will likely be associated with individuals resting on the sea during the hours of darkness in pelagic habitats; although skuas may also migrate at night, especially during full moons (Bonnet-Lebrun et al. 2021). However, this also provides evidence that the skuas incorporated a fly-and-forage migration strategy (Strandberg & Alerstam 2007, Amélineau et al. 2021).

### Population specific variation in migration strategies

We found little variation in the migration strategies of skuas from Scotland, Faroe Islands and Norway. However, there was a population difference in time spent at stopovers driven by individuals from Svalbard, the most northerly tracked breeding population, which spent a lower proportion of their northbound migration at stopovers. Arctic Skuas from Svalbard have temporally shorter northbound migrations and spend less time in the North Atlantic compared to skuas from other populations as well as arriving back to their breeding areas later (van Bemmelen et al. 2023). This may indicate that Arctic Skuas from Svalbard either fuel their migration to a larger extent with reserves acquired at the wintering areas, or attain higher daily travel speeds due to more efficient or longer time spent foraging per day. The latter would be facilitated by longer day-lengths later in the spring. This variation in migration strategies may in part account for the different population trends of Arctic Skuas in the North Atlantic, with the Svalbard breeding population declining less than the other populations. However, the mechanisms that might link migratory strategies to the skuas population status are unknown emphasizing the need to better understand the carry-over effects of different migration strategies on subsequent breeding success and survival (Finch et al. 2014, Szostek & Becker 2015, Bogdanova et al. 2017).

Where the skuas wintered also influenced their migration strategies. Despite the greater distances and energetic costs involved, skuas wintering in the most southern areas, around the Benguela Current and Patagonian Shelf, typically spent a lower proportion of their migrations at stopovers than those migrating to closer wintering areas. This indicates that instead of spending additional time at stopovers to fuel their longer migrations, the skuas employed a fly-and-forage migration strategy. Here, individuals combine opportunistic foraging with directional transit flights, which has been observed in numerous seabird species, especially those with a low wing loading and lower cost of flight, such as Arctic Skuas (Strandberg & Alerstam 2007, Alerstam & Bäckman 2018, Amélineau et al. 2021). This strategy should particularly benefit individuals migrating to the furthest wintering areas as they can increase the speed of their migration whilst maintaining energy requirements. Although core stopovers made by skuas coincided with areas associated with predictable, high marine productivity, some stopovers also occurred outside these commonly used staging areas, further indicating that individuals foraged opportunistically along their migration routes. Undertaking a fly-and-forage strategy, in combination with periods in staging areas, may buffer individuals from potential adverse environmental conditions to a greater extent than those which rely entirely on staging areas to refuel. Conversely, individuals that are reliant on finding sufficient food as they migrate maybe at a disadvantage to those that rely largely on staging areas, especially with depleting marine resources (Cury et al. 2011, Grémillet et al. 2018), which may also contribute to the observed Arctic Skua declines. Therefore, where a fly-and-forage strategy is key to successful migration it is important that habitats that provide foraging opportunities along migratory routes are also protected (Amélineau et al. 2021, Rueda-Uribe et al. 2022), for the skuas and the species they may kleptoparasitise on route (Egevang et al. 2010, Stenhouse et al. 2012).

## Conclusion

This study reveals the importance of several staging areas of high marine productivity that Arctic Skuas used consistently for refuelling across years and colonies. However, it also highlights the potential vulnerability of populations to stressors that might occur in these discrete locations, for example, associated with severe weather events and competition with fisheries. Identifying key staging areas provides important evidence for the designation of marine protected areas outside the breeding season (Davies et al. 2021), specifically the staging area off West Africa, which is not currently designated, and was not previously known to be important for migrating Arctic Skuas. Differences in migration routes may also result in differential risk to specific stressors, such as the potential risks associated with offshore renewable developments in the North Sea. Having better data on where Arctic Skuas are located outside the breeding season means that overlap between at-sea distributions and potential threats can be used to assess and prioritize risk and associated mitigation measures (Phillips et al. 2016, Hays et al. 2019). Implementing large-scale actions such as improving fisheries regulations to better manage fish stocks, and reduce disturbance and mortality from bycatch and ship strikes, would also benefit long-distance migrants, both within staging areas and along migration routes (Gremillet et al. 2015, Oppel et al. 2019).

By identifying population differences in migration routes and strategies we can also better investigate the drivers of differential population trends. It would be advantageous to explore whether the slightly different migration strategies and staging areas of Svalbard individuals have positive carry-over effects to subsequent breeding seasons. By establishing multi-national long-term demographic studies across the range of species, we can better link population differences in migration strategies, routes and staging areas to specific aspects of demography (Harrison et al 2021). Integration of demography and phenology with tracking data can also help establish whether differences in migration connectivity and strategies occur among different demographic groups, for example associated with sex or age, which may have consequences on populations dynamics if specific groups experience different levels of risk during migration (Briedis & Bauer 2018, Aldará et al. 2019).

Identifying where long-distance migrants, such as Arctic Skuas, are distributed during migration and the strategies they undertake is a vital first step in determining the stressors individuals may encounter on route, and linking carry-over effects to demography and population dynamics; all of which are necessary to understand population declines and prioritise conservation actions under continued environmental change (Martin et al. 2007, Sheehy et al. 2010, Strøm et al. 2021, Santos et al. in prep).

## Acknowledgements

Special thanks to everyone who helped with fieldwork and all landowners. In Slettnes, we were assisted in the field by Geert Aarts, Daniël van Denderen, Jan van Dijk, Maria van Leeuwe, Daan Liefhebber, Morrison Pot, Marc van Roomen, Janne Schekkerman and Rinse van der Vliet. Other help was provided by Cees and Mimi Tesselaar, Barbara Ganther, Hans-Ulrich Rösner, Karl-Birger Strann, Jeanette Hickman, Torstein Johnsrud, Niels Westpahl and Benedicte Færevåg. In Svalbard and Brensholmen, help was provided by Erlend Lorentzen, Elise Biersma, Fokje Schaafsma, Anouk Goedknegt, Oebele Dijk, Maarten Loonen, Anette Fenstad, Elise Skottene, Liv Monica Trondrud, Nora Bjørnlid, Heidi Kilen, Eline Rypdal, Emilly Hill, Melissa Fontenille and Thomas Oudman. In the Faroe Islands, help was provided by Jón Aldará, Leivur Janus Hansen and Anthony Weatherhill; we kindly acknowledge the landowners on Fugloy for permission to work on their property. Support in the field in Scotland was provided by Fair Isle Bird Observatory and NatureScot. James Fox (Migrate Technology Ltd) and Glen Fowler (Biotrack Ltd) kindly retrieved data from loggers that failed to download. BTO research was funded by several generous individual donors. The work at Slettnes was financed by the Netherlands Organisation for Scientific Research (NWO project number 866.13.005) and supported by The Norwegian Institute for Nature Research (NINA). Thanks to the Faroese Research Council which supported the Faroese fieldwork financially. Catching and deploying geolocators was approved in Scotland by the Special Method Technical Panel, part of the BTO/JNCC Avian Demographic Scheme; in the Faroes Islands by the Copenhagen Bird Ringing Centre and The National Museum of the Faroe Islands; and in Svalbard and Norway by the Governor of Svalbard and Norwegian Food Safety Authority (FOTS ID 2086, 3817, 6329, 8538, 15726).

## Appendix. Similarity of migration routes of individuals among years

For 46 individuals across breeding populations, we had data on full migratory routes for multiple years. Therefore, to establish whether routes were more similar within-(pairs of tracks completed by the same individuals) than 1) those between-individuals with the same breeding population and wintering areas, and 2) those between-individuals from different breeding populations and/or wintering locations we calculated dynamic time warping (DTW) ratios. DTW can be used to quantify the similarity of migration routes of different lengths that can vary in time and speed (Senin 2008, Ranacher & Tzavella 2014, Cleasby et al. 2019), as did our Arctic Skuas tracks. DTW calculates the unbounded minimal cumulative distance path between two tracks, with larger values indicating larger distances between pairs of tracks, and which are therefore more dissimilar (Cleasby et al. 2019). As the skua tracks differed in length depending on the start and finish locations, as well as the length of gaps resulting from the equinoxes, we calculated DTW ratios by dividing the DTW value of each pair of tracks by the length of the larger track, to prevent track length overly influencing the results (Cleasby et al. 2019). Pairwise DTW ratios were calculated for double smoothed tracks, of at least ten positions, for southbound and northbound separately, using the *SimilarityMeasures* R package (Toohey 2015).

To test for individual consistency in migratory routes, across populations we performed Kruskal-Wallis rank sum tests, for southbound and northbound separately, between the DTW ratio and whether pairs of routes were within- or between-individuals. With between-individuals being split into two groups: 1) those with the same breeding population and wintering areas, and 2) those from different breeding populations and/or wintering locations.

Based on the DTW ratios, during southbound migration, routes were significantly more similar within-individuals (10.8 ± SD 8.2, 112 paired tracks from 46 individuals) than between-individuals from the same population and wintering location (19.4 ± 17.1, 2006 paired tracks from 95 individuals), and between-individuals from different populations and/or wintering locations (28.7 ± 16.6, 14353 paired tracks from 109 individuals) across all years (Kruskal-Wallis test, *H* = 984.9, df = 2, *P* < 0.001). This was also the case during northbound migration, with routes significantly more similar within-individuals (6.7 ± SD 3.4, 97 paired tracks from 45 individuals) than between-individuals from the same population and wintering location (14.0 ± 8.8, 1585 paired tracks from 96 individuals), and between-individuals from different populations and/or wintering locations (22.4 ± 12.1, 13894 paired tracks from 109 individuals) across all years (Kruskal-Wallis test, *H* = 1085.5, df = 2, *P* < 0.001).

## Notes

### Competing Interest Statement

The authors have declared no competing interest.

## References

1. Aldará J, Hammer S, Thorup K, Snell KRS (2019) Determining hatch dates for skuas: an egg density calibration curve. Seabird 32:84–95.

2. Alerstam T, Bäckman J. (2018) Ecology of animal migration. Curr Biol 28: R968–R972.

3. Ambrosini R, Møller AP, Saino N (2009) A quantitative measure of migratory connectivity. Journal of Theoretical Biology 257: 203–211.

4. Amélineau F, Merkel B, Tarroux A, Descamps S, Anker-Nilssen T, Bjørnstad O, Bråthen V, Chastel O, Christensen-Dalsgaard S, Danielsen J, Daunt F, Dehnhard N, Ekker M, Erikstad K, Ezhov A, Fauchald P, Gavrilo M, Hallgrimsson G, Hansen E, Harris M, Helberg M, Helgason H, Johansen M, Jónsson J, Kolbeinsson Y, Krasnov Y, Langset M, Lorentsen S, Lorentzen E, Melnikov M, Moe B, Newell M, Olsen B, Reiertsen T, Systad G, Thompson P, Thórarinsson T, Tolmacheva E, Wanless S, Wojczulanis-Jakubas K, Åström J, Strøm H (2021) Six pelagic seabird species of the North Atlantic engage in a fly-and-forage strategy during their migratory movements. Mar Ecol Prog Ser 676:127–144.

5. Bauer S, Lisovski S, Hahn S (2016) Timing is crucial for consequences of migratory connectivity. Oikos 125:605–612.

6. Belisle M, Giroux JF (1995) Predation and kleptoparasitism by migrating parasitic jaegers. Condor 97:771–781.

7. Belopollskiil LO (1961). Ecology of sea colony birds of the Barents Sea.

8. van Bemmelen RSA, Schekkerman H, Hin V, Pot MT, Janssen K, Ganter B, Rösner HU, Tulp I (2021) Heavy decline of the largest European Arctic Skua *Stercorarius parasiticus* colony: interacting effects of food shortage and predation. Bird Study 68:100–111.

9. van Bemmelen RSA, Moe B, Schekkerman H, Hansen SA, Snell KRS, Humphreys EM, Mantyla E, Hallgrimsson GT, Gilg O, Ehrich D, Calladine J, Hammer S, Harris SJ, Lang J, Vignisson SR, Kolbeinsson Y, Nuotio K, Sillanpaa M, Sittler B, Sokolov A, Klaassen RHG, Phillips RA, Tulp I (2023) Ocean scale variation in migration schedules of a long distance migratory seabird is fully compensated upon return to the breeding site. biorxiv 2023.05.27.542544.

10. Bhattacharyya A (1943) On a measure of divergence between two statistical populations defined by their probability distributions. Bull Calcutta Math Soc 35:99–110.

11. BirdLife International (2021) IUCN Red List for Birds. http://datazone.birdlife.org/species/search (accessed 18 March 2021)

12. Bivand R, Rundel C (2019) Rgeos: Interface to Geometry Engine - Open Source (’GEOS’). R package version 0.4.

13. Bogdanova MI, Butler A, Wanless S, Moe B, Anker-Nilssen T, Frederiksen M, Boulinier T, Chivers LS, Christensen-Dalsgaard S, Descamps S, Harris MP, Newell M, Olsen B, Phillips RA, Shaw D, Steen H, Strøm H, Thórarinsson TL, Daunt F (2017) Multi-colony tracking reveals spatio-temporal variation in carry-over effects between breeding success and winter movements in a pelagic seabird. Mar Ecol Prog Ser 578:167–181.

14. Bonnet-Lebrun A-S, Dias MP, Phillips RA, Granadeiro JP, Brooke M de L, Chastel O, Clay TA, Fayet AL, Gilg O, González-Solís J, Guilford T, Hanssen SA, Hedd A, Jaeger A, Krietsch J, Lang J, Le Corre M, Militão T, Moe B, Montevecchi WA, Peter H-U, Pinet P, Rayner MJ, Reid T, Reyes-González JM, Ryan PG, Sagar PM, Schmidt NM, Thompson DR, van Bemmelen R, Watanuki Y, Weimerskirch H, Yamamoto T, Catry P (2021) Seabird migration strategies: Flight budgets, diel activity patterns, and lunar influence. Front Mar Sci 8:1–15.

15. Briedis M, Bauer S (2018) Migratory connectivity in the context of differential migration. Biol Lett 14: 20180679.

16. Brooks ME, Kristensen K, van Benthem KJ, Magnusson A, Berg CW, Nielsen A, Skaug HJ, Mächler M, Bolker BM (2017) GlmmTMB balances speed and flexibility among packages for zero-inflated generalized linear mixed modeling. R J 9:378–400.

17. Brown JM, van Loon EE, Bouten W, Camphuysen KCJ, Lens L, Müller W, Thaxter CB, Shamoun-Baranes J (2021) Long-distance migrants vary migratory behaviour as much as short-distance migrants: An individual-level comparison from a seabird species with diverse migration strategies. J Anim Ecol 90:1058–1070.

18. Buechley ER, Oppel S, Efrat R, Phipps WL, Carbonell Alanís I, Álvarez E, Andreotti A, Arkumarev V, Berger-Tal O, Bermejo Bermejo A, Bounas A, Ceccolini G, Cenerini A, Dobrev V, Duriez O, García J, García-Ripollés C, Galán M, Gil A, Giraud L, Hatzofe O, Iglesias-Lebrija JJ, Karyakin I, Kobierzycki E, Kret E, Loercher F, López-López P, Miller Y, Mueller T, Nikolov SC, de la Puente J, Sapir N, Saravia V, Şekercioğlu ÇH, Sillett TS, Tavares J, Urios V, Marra PP (2021) Differential survival throughout the full annual cycle of a migratory bird presents a life-history trade-off. J Anim Ecol 90:1228–1238.

19. Burger AE, Shaffer SA (2008) Application of tracking and data-logging technology in research and conservation of seabirds. Auk 125:253–264.

20. Calenge C (2006) The package “adehabitat” for the R software: A tool for the analysis of space and habitat use by animals. Ecol Modell 197:516–519.

21. Calenge C (2015) Package ‘adehabitat’. R package version, 1, 18.

22. Caliendo V, Lewis NS, Pohlmann A, Baillie SR, Banyard AC (2022) Transatlantic spread of highly pathogenic avian influenza H5N1 by wild birds from Europe to North America in 2021. Sci Rep:1–18.

23. Camphuysen CJ, Van Der Meer J (2005) Wintering seabirds in West Africa: Foraging hotspots off Western Sahara and Mauritania driven by upwelling and fisheries. African J Mar Sci 27:427–437.

24. Campioni L, Dias MP, Granadeiro JP, Catry P (2020) An ontogenetic perspective on migratory strategy of a long-lived pelagic seabird: Timings and destinations change progressively during maturation. J Anim Ecol 89:29–43.

25. Cashion T, Nguyen T, Brink T ten, Mook A, Palacios-Abrantes J, Roberts SM (2020) Shifting seas, shifting boundaries: Dynamic marine protected area designs for a changing climate. PLoS One 15:1–17.

26. Chirosca AM, Rusu L, Bleoju A (2022) Study on wind farms in the North Sea area. Energy Reports 8:162–168.

27. Cohen EB, Hostetler JA, Hallworth MT, Rushing CS, Sillett TS, Marra PP (2017) Quantifying the strength of migratory connectivity. Methods Ecol Evol 9:513–524.

28. Cohen EB, Rushing CR, Moore FR, Hallworth MT, Hostetler JA, Ramirez MG, Marra PP (2018) The strength of migratory connectivity for birds en route to breeding through the Gulf of Mexico. Ecography (Cop) 42:658–669.

29. Conklin JR, Battley PF, Potter MA, Fox JW (2010) Breeding latitude drives individual schedules in a trans-hemispheric migrant bird. Nat Commun 1:3–8.

30. Cury PM, Boyd IL, Bonhommeau S, Anker-Nilssen T, Crawford RJM, Furness RW, Mills JA, Murphy EJ, Osterblom H, Paleczny M, Piatt JF, Roux J-P, Shannon L, Sydeman WJ (2011) Global seabird response to forage fish depletion-one-third for the birds. Science 334:1703–6.

31. Davies TE, Carneiro APB, Tarzia M, Wakefield E, Hennicke JC, Frederiksen M, Hansen ES, Campos B, Hazin C, Lascelles B, Anker-Nilssen T, Arnardóttir H, Barrett RT, Biscoito M, Bollache L, Boulinier T, Catry P, Ceia FR, Chastel O, Christensen-Dalsgaard S, Cruz-Flores M, Danielsen J, Daunt F, Dunn E, Egevang C, Fagundes AI, Fayet AL, Fort J, Furness RW, Gilg O, González-Solís J, Granadeiro JP, Grémillet D, Guilford T, Hanssen SA, Harris MP, Hedd A, Huffeldt NP, Jessopp M, Kolbeinsson Y, Krietsch J, Lang J, Linnebjerg JF, Lorentsen S, Madeiros J, Magnusdottir E, Mallory ML, McFarlane Tranquilla L, Merkel FR, Militão T, Moe B, Montevecchi WA, Morera-Pujol V, Mosbech A, Neves V, Newell MA, Olsen B, Paiva VH, Peter H, Petersen A, Phillips RA, Ramírez I, Ramos JA, Ramos R, Ronconi RA, Ryan PG, Schmidt NM, Sigurðsson IA, Sittler B, Steen H, Stenhouse IJ, Strøm H, Systad GHR, Thompson P, Thórarinsson TL, van Bemmelen RSA, Wanless S, Zino F, Dias MP (2021) Multispecies tracking reveals a major seabird hotspot in the North Atlantic. Conserv Lett e12824.

32. Desprez M, Jenouvrier S, Barbraud C, Delord K, Weimerskirch H (2018) Linking oceanographic conditions, migratory schedules and foraging behaviour during the non-breeding season to reproductive performance in a long-lived seabird. Funct Ecol 32:2040–2053.

33. Dias MP, Granadeiro JP, Catry P (2012) Do seabirds differ from other migrants in their travel arrangements? On route strategies of Cory’s Shearwater during its trans-equatorial journey. PLoS One 7:e49376.

34. Dias MP, Martin R, Pearmain EJ, Burfield IJ, Small C, Phillips RA, Yates O, Lascelles B, Borboroglu PG, Croxall JP (2019) Threats to seabirds: A global assessment. Biol Conserv 237:525–537.

35. Dunn DC, Harrison AL, Curtice C, DeLand S, Donnelly B, Fujioka E, Heywood E, Kot CY, Poulin S, Whitten M, Åkesson S, Alberini A, Appeltans W, Arcos JM, Bailey H, Ballance LT, Block B, Blondin H, Boustany AM, Brenner J, Catry P, Cejudo D, Cleary J, Corkeron P, Costa DP, Coyne M, Crespo GO, Davies TE, Dias MP, Douvere F, Ferretti F, Formia A, Freestone D, Friedlaender AS, Frisch-Nwakanma H, Froján CB, Gjerde KM, Glowka L, Godley BJ, Gonzalez-Solis J, Granadeiro JP, Gunn V, Hashimoto Y, Hawkes LM, Hays GC, Hazin C, Jimenez J, Johnson DE, Luschi P, Maxwell SM, McClellan C, Modest M, Di Sciara GN, Palacio AH, Palacios DM, Pauly A, Rayner M, Rees AF, Salazar ER, Secor D, Sequeira AMM, Spalding M, Spina F, Van Parijs S, Wallace B, Varo-Cruz N, Virtue M, Weimerskirch H, Wilson L, Woodward B, Halpin PN (2019) The importance of migratory connectivity for global ocean policy. Proc R Soc B Biol Sci 286:20191472.

36. Egevang C, Stenhouse IJ, Phillips RA, Petersen A, Fox JW, Silk JRD (2010) Tracking of Arctic terns Sterna paradisaea reveals longest animal migration. Proc Natl Acad Sci U S A 107:2078–2081.

37. Fayet AL, Freeman R, Anker-Nilssen T, Diamond A, Erikstad KE, Fifield D, Fitzsimmons MG, Hansen ES, Harris MP, Jessopp M, Kouwenberg AL, Kress S, Mowat S, Perrins CM, Petersen A, Petersen IK, Reiertsen TK, Robertson GJ, Shannon P, Sigurðsson IA, Shoji A, Wanless S, Guilford T (2017) Ocean-wide drivers of migration strategies and their influence on population breeding performance in a declining seabird. Curr Biol 27:3871–3878.e3.

38. Fieberg J, Kochanny CO (2005) Quantifying home-range overlap: the importance of the utilization distribution. J Wildl Manage 69:1346–1359.

39. Finch T, Pearce-Higgins JW, Leech DI, Evans KL (2014) Carry-over effects from passage regions are more important than breeding climate in determining the breeding phenology and performance of three avian migrants of conservation concern. Biodivers Conserv 23:2427–2444.

40. Frederiksen M, Moe B, Daunt F, Phillips RA, Barrett RT, Bogdanova MI, Boulinier T, Chardine JW, Chastel O, Chivers LS, Christensen-Dalsgaard S, Clément-Chastel C, Colhoun K, Freeman R, Gaston AJ, González-Solís J, Goutte A, Grémillet D, Guilford T, Jensen GH, Krasnov Y, Lorentsen SH, Mallory ML, Newell M, Olsen B, Shaw D, Steen H, Strøm H, Systad GH, Thórarinsson TL, Anker-Nilssen T (2012) Multicolony tracking reveals the winter distribution of a pelagic seabird on an ocean basin scale. Divers Distrib 18:530–542.

41. Fretwell D (1972) Populations in a Seasonal Environment. Princeton University Press.

42. Furness RW (1987) The skuas. T and AD Poyser, Calton, England.

43. Furness RW, Hallgrimsson GT, Montevecchi WA, Fifield D, Kubetzki U, Mendel B, Garthe S (2018) Adult Gannet migrations frequently loop clockwise around Britain and Ireland. Ringing Migr 33:45–53.

44. Furness RW, Wade HM, Masden EA (2013) Assessing vulnerability of marine bird populations to offshore wind farms. J Environ Manage 119:56–66.

45. Gilg O, Moe B, Hanssen SA, Schmidt NM, Sittler B, Hansen J, Reneerkens J, Sabard B, Chastel O, Moreau J, Phillips RA, Oudman T, Biersma EM, Fenstad AA, Lang J, Bollache L (2013) Trans-equatorial migration routes, staging sites and wintering areas of a high-Arctic avian predator: The Long-tailed Skua (*Stercorarius longicaudus*). PLoS One 8:e64614.

46. Gilroy JJ, Gill JA, Butchart SHM, Jones VR, Franco AMA (2016) Migratory diversity predicts population declines in birds. Ecol Lett 19:308–317.

47. Grecian WJ, Witt MJ, Attrill MJ, Bearhop S, Becker PH, Egevang C, Furness RW, Godley BJ, González-Solís J, Grémillet D, Kopp M, Lescroël A, Matthiopoulos J, Patrick SC, Peter HU, Phillips RA, Stenhouse IJ, Votier SC (2016) Seabird diversity hotspot linked to ocean productivity in the Canary Current Large Marine Ecosystem. Biol Lett 12:20160024.

48. Grémillet D, Boulinier T (2009) Spatial ecology and conservation of seabirds facing global climate change: a review. Mar Ecol Prog Ser 391:121–137.

49. Gremillet D, Peron C, Provost P, Lescroel A (2015) Adult and juvenile European seabirds at risk from marine plundering off West Africa. Biol Conserv 182:143–147.

50. Grémillet D, Ponchon A, Paleczny M, Palomares MLD, Karpouzi V, Pauly D (2018) Persisting worldwide seabird-fishery competition despite seabird community decline. Curr Biol 28:4009–4013.e2.

51. Harrison L, Woodard PF, Mallory ML, Rausch J, Harrison L, Bird M (2021) Sympatrically breeding congeneric seabirds (*Stercorarius* spp.) from Arctic Canada migrate to four oceans. Ecol Evol 00:1–12.

52. Harrison XA, Blount JD, Inger R, Norris DR, Bearhop S (2011) Carry-over effects as drivers of fitness differences in animals. J Anim Ecol 80:4–18.

53. Hays GC, Bailey H, Bograd SJ, Bowen WD, Campagna C, Carmichael RH, Casale P, Chiaradia A, Costa DP, Cuevas E, Nico de Bruyn PJ, Dias MP, Duarte CM, Dunn DC, Dutton PH, Esteban N, Friedlaender A, Goetz KT, Godley BJ, Halpin PN, Hamann M, Hammerschlag N, Harcourt R, Harrison AL, Hazen EL, Heupel MR, Hoyt E, Humphries NE, Kot CY, Lea JSE, Marsh H, Maxwell SM, McMahon CR, Notarbartolo di Sciara G, Palacios DM, Phillips RA, Righton D, Schofield G, Seminoff JA, Simpfendorfer CA, Sims DW, Takahashi A, Tetley MJ, Thums M, Trathan PN, Villegas-Amtmann S, Wells RS, Whiting SD, Wildermann NE, Sequeira AMM (2019) Translating marine animal tracking data into conservation policy and management. Trends Ecol Evol 34:459–473.

54. Henriksen S, Hilmo O (2015) Norsk rødliste for arter 2015. Artsdatabanken, Norge ISBN: 978–82-92838-40-2

55. Hijmans R (2019) Geosphere: Spherical Trigonometry. R package version 1.5–10.

56. Knight EC, Harrison A-L, Scarpignato AL, Van Wilgenburg SL, Bayne EM, Ng JW, Angell E, Bowman R, Brigham RM, Drolet B, Easton WE, Forrester TR, Foster JT, Haché S, Hannah KC, Tremblay JA, Marra PP (2021) Comprehensive estimation of spatial and temporal migratory connectivity across the annual cycle to direct conservation efforts. Ecograph 44:665–679.

57. Knudsen E, Lindén A, Both C, Jonzén N, Pulido F, Saino N, Sutherland WJ, Bach LA, Coppack T, Ergon T, Gienapp P, Gill JA, Gordo O, Hedenström A, Lehikoinen E, Marra PP, Møller AP, Nilsson ALK, Péron G, Ranta E, Rubolini D, Sparks TH, Spina F, Studds CE, Sæther SA, Tryjanowski P, Stenseth NC (2011) Challenging claims in the study of migratory birds and climate change. Biol Rev 86:928–946.

58. Koenig WD, Liebhold AM (2016) Temporally increasing spatial synchrony of North American temperature and bird populations. Nat Clim Chang 6:614–617.

59. Kubelka V, Sandercock BK, Székely T, Freckleton RP (2021) Animal migration to northern latitudes: environmental changes and increasing threats. Trends Ecol Evol:1–12.

60. Lamb JS, Satgé YG, Jodice PGR (2017) Influence of density-dependent competition on foraging and migratory behavior of a subtropical colonial seabird. Ecol Evol 7:6469– 6481.

61. Lascelles BG, Taylor PR, Miller MGR, Dias MP, Oppel S, Torres L, Hedd A, Le Corre M, Phillips RA, Shaffer SA, Weimerskirch H, Small C (2016) Applying global criteria to tracking data to define important areas for marine conservation. Divers Distrib 22:422– 431.

62. Lecomte VJ, Sorci G, Cornet S, Jaeger A, Faivre B, Arnoux E, Gaillard M, Trouvé C, Besson D, Chastel O, Weimerskirch H (2010) Patterns of aging in the long-lived wandering albatross. Proc Natl Acad Sci U S A 107:6370–6375.

63. Lenth VR (2021) emmeans: Estimated Marginal Means, aka Least-Squares Means. R package version 1.7.0. https://CRAN.R-project.org/package=emmeans

64. Lisovski S, Hahn S (2012) GeoLight - processing and analysing light-based geolocator data in R. Methods Ecol Evol 3:1055–1059.

65. Marra P, Hobson KA, Holmes RT (1998) Linking winter and summer events in a migratory bird by using stable-carbon isotopes. Science 282:1884–1886.

66. Marra PP, Cohen EB, Loss SR, Rutter JE, Tonra CM (2015) A call for full annual cycle research in animal ecology. Biol Lett 11:2015.0552.

67. Martin TG, Chadès I, Arcese P, Marra PP, Possingham HP, Norris DR (2007) Optimal conservation of migratory species. PLoS One 2:3–7.

68. McClintock BT, Michelot T (2021) MomentuHMM: R package for generalized hidden Markov models of animal movement. Methods Ecol Evol 9:1518–1530.

69. Merkel B, Descamps S, Yoccoz N, Grémillet D, Fauchald P, Danielsen J, Daunt F, Erikstad K, Ezhov A, Harris M, Gavrilo M, Lorentsen S, Reiertsen T, Systad G, Lindberg Thórarinsson T, Wanless S, Strøm H (2021) Strong migratory connectivity across meta-populations of sympatric North Atlantic seabirds. Mar Ecol Prog Ser 676:173–188.

70. Newton I (2008) The Migration Ecology of Birds. Academic Press.

71. Oppel S, Bolton M, Carneiro APB, Dias MP, Green JA, Masello JF, Phillips RA, Owen E, Quillfeldt P, Beard A, Bertrand S, Blackburn J, Boersma PD, Borges A, Broderick AC, Catry P, Cleasby I, Clingham E, Creuwels J, Crofts S, Cuthbert RJ, Dallmeijer H, Davies D, Davies R, Dilley BJ, Dinis HA, Dossa J, Dunn MJ, Efe MA, Fayet AL, Figueiredo L, Frederico AP, Gjerdrum C, Godley BJ, Granadeiro JP, Guilford T, Hamer KC, Hazin C, Hedd A, Henry L, Hernández-Montero M, Hinke J, Kokubun N, Leat E, Tranquilla LMF, Metzger B, Militão T, Montrond G, Mullié W, Padget O, Pearmain EJ, Pollet IL, Pütz K, Quintana F, Ratcliffe N, Ronconi RA, Ryan PG, Saldanha S, Shoji A, Sim J, Small C, Soanes L, Takahashi A, Trathan P, Trivelpiece W, Veen J, Wakefield E, Weber N, Weber S, Zango L, González-Solís J, Croxall J (2018) Spatial scales of marine conservation management for breeding seabirds. Mar Policy 98:37–46.

72. Palm EC, Newman SH, Prosser DJ, Xiao X, Ze L, Batbayar N, Balachandran S, Takekawa JY (2015) Mapping migratory flyways in Asia using dynamic Brownian bridge movement models. Mov Ecol 3:1–10.

73. Paterson AM (1986) Kleptoparasitic feeding by migrating skuas in Malaga Bay, Spain. Ringing Migr 7:51–55.

74. Pelletier D, Seyer Y, Garthe S, Bonnefoi S, Phillips RA, Guillemette M (2020) So far, so good. . . Similar fitness consequences and overall energetic costs for short and long-distance migrants in a seabird. PLoS One 15:1–23.

75. Perkins A, Ratcliffe N, Suddaby D, Ribbands B, Smith C, Ellis P, Meek E, Bolton M (2018) Combined bottom-up and top-down pressures drive catastrophic population declines of Arctic skuas in Scotland. J Anim Ecol 87:1573–1586.

76. Phillips RA, Gales R, Baker GB, Double MC, Favero M, Quintana F, Tasker ML, Weimerskirch H, Uhart M, Wolfaardt A (2016) The conservation status and priorities for albatrosses and large petrels. Biol Conserv 201:169–183.

77. Phillips RA, Silk JRD, Croxall JP, Afanasyev V, Briggs DR (2004) Accuracy of geolocation estimates for flying seabirds. Mar Ecol Prog Ser 266:265–272.

78. R Core Development Team (2021) R: A language and environment for statistical computing.

79. Rakhimberdiev E, Duijns S, Karagicheva J, Camphuysen CJ, Castricum VRS, Dekinga A, Dekker R, Gavrilov A, Horn J, Jukema J, Saveliev A, Soloviev M, Tibbitts TL, Gils JA Van, Piersma T (2018) Fuelling conditions at staging sites can mitigate Arctic warming effects in a migratory bird. Nat Commun 9:4263.

80. Ramírez F, Afán I, Davis LS, Chiaradia A (2017) Climate impacts on global hot spots of marine biodiversity. Sci Adv 3:1–8.

81. Reiertsen T, Layton-Matthews K, Erikstad K, Hodges K, Ballesteros M, Anker-Nilssen T, Barrett R, Benjaminsen S, Bogdanova M, Christensen-Dalsgaard S, Daunt F, Dehnhard N, Harris M, Langset M, Lorentsen S, Newell M, Bråthen V, Støyle-Bringsvor I, Systad G, Wanless S (2021) Inter-population synchrony in adult survival and effects of climate and extreme weather in non-breeding areas of Atlantic puffins. Mar Ecol Prog Ser 676:219–231.

82. Rueda-Uribe C, Lötberg U, Åkesson S (2022) Foraging on the wing for fish while migrating over changing landscapes: traveling behaviors vary with available aquatic habitat for Caspian terns. Mov Ecol 10:1–15.

83. Rushing CS, Ryder TB, Marra PP (2016) Quantifying drivers of population dynamics for a migratory bird throughout the annual cycle. Proc R Soc B Biol Sci 283:20152846.

84. Santos I (2018) Survival and breeding success of the declining Arctic Skua population of the Faroe Islands. University of Copenhagen, Denmark

85. Santos IAM dos, Snell KRS, van Bemmelen RS, Moe B, Thorup K (in prep.) Wintering, rather than reeding, oceanic conditions contribute to declining survival in a long-distance migratory seabird.

86. Schultner J, Moe B, Chastel O, Bech C, Kitaysky AS (2014) Migration and stress during reproduction govern telomere dynamics in a seabird. Biol Lett 10.

87. Seyer Y, Gauthier G, Bêty J, Therrien J, Lecomte N (2021) Seasonal variations in migration strategy of a long-distance arctic-breeding seabird. Mar Ecol Prog Ser 677:1–16.

88. Sheehy J, Taylor CM, Mccann KS, Norris DR (2010) Optimal conservation planning for migratory animals: Integrating demographic information across seasons. Conserv Lett 3:192–202.

89. Stenhouse IJ, Egevang C, Phillips RA (2012) Trans-equatorial migration, staging sites and wintering area of Sabine’s Gulls *Larus sabini* in the Atlantic Ocean. Ibis 154:42–51.

90. Strandberg R, Alerstam T (2007) The strategy of fly-and-forage migration, illustrated for the osprey (*Pandion haliaetus*). Behav Ecol Sociobiol 61:1865–1875.

91. Strandberg R, Klaassen RHG, Thorup K (2009) Spatio-temporal distribution of migrating raptors: a comparison of ringing and satellite tracking. J Avian Biol 40:500–510.

92. Strøm H, Descamps S, Ekker M, Fauchald P, Moe B (2021) Tracking the movements of North Atlantic seabirds: steps towards a better understanding of population dynamics and marine ecosystem conservation. Mar Ecol Prog Ser 676:97–116.

93. Szostek KL, Becker PH (2015) Survival and local recruitment are driven by environmental carry-over effects from the wintering area in a migratory seabird. Oecologia:643–657.

94. Trierweiler C, Klaassen RHG, Drent RH, Exo KM, Komdeur J, Bairlein F, Koks BJ (2014) Migratory connectivity and population specific migration routes in a long-distance migratory bird. Proc R Soc B Biol Sci 281: 20132897.

95. Trierweiler C, Mullié WC, Drent RH, Exo KM, Komdeur J, Bairlein F, Harouna A, De Bakker M, Koks BJ (2013) A Palaearctic migratory raptor species tracks shifting prey availability within its wintering range in the Sahel. J Anim Ecol 82:107–120.

96. Tuck GN, Polacheck T, Croxall JP, Weimerskirch H, Prince PA, Wotherspoon S (1999) The potential of archival tags to provide long-term movement and behaviour data for seabirds: First results from Wandering Albatross *Diomedea exulans* of South Georgia and the Crozet Islands. Emu 99:60–68.

97. Warnock N (2010) Stopping vs. staging: The difference between a hop and a jump. J Avian Biol 41:621–626.

98. Wickham H (2016) Ggplot2: Elegant Graphics for Data Analysis. Springer-Verlag New York.

99. Wilson RP, Grémillet D, Syder J, Kierspel MAM, Garthe S, Weimerskirch H, Schäfer-neth C, Scolaro JA, Bost C, Plötz J, Nel D (2002) Remote-sensing systems and seabirds: their use, abuse and potential for measuring marine environmental variables. Mar Ecol Prog Ser 228:241–261.

100. Wuorinen JD (1992) Do Arctic Skuas *Stercorarius parasiticus* exploit and follow terns during the fall migration? Ornis Fenn 69:198–200.

## References

102. Cleasby IR, Wakefield ED, Morrissey BJ, Bodey TW, Votier SC, Bearhop S, Hamer KC (2019) Using time-series similarity measures to compare animal movement trajectories in ecology. Behav Ecol Sociobiol 73:151.

103. Ranacher P, Tzavella K (2014) How to compare movement? A review of physical movement similarity measures in geographic information science and beyond. Cartogr Geogr Inf Sci 41:286–307.

104. Senin P (2008) Dynamic Time Warping Algorithm Review. Manoa Honolulu, USA.

