## Supplementary Material for "Cross population comparison of complex migration strategies in a declining oceanic seabird"

**Table S1.** Number of Arctic Skuas tracks by population and year. ‘Years tracked’ represent the time between two breeding periods.

| Colonies | Population | Years tracked | No. of tracks |
| --- | --- | --- | --- |
| Fugloy | Faroe Islands | 2016/17 | 21 |
| Fugloy | Faroe Islands | 2017/18 | 6 |
| Brensholmen and Slettnes | Norway | 2011/12 | 6 |
| Brensholmen and Slettnes | Norway | 2012/13 | 6 |
| Brensholmen and Slettnes | Norway | 2013/14 | 3 |
| Brensholmen and Slettnes | Norway | 2014/15 | 27 |
| Brensholmen and Slettnes | Norway | 2015/16 | 28 |
| Brensholmen and Slettnes | Norway | 2016/17 | 16 |
| Brensholmen and Slettnes | Norway | 2017/18 | 13 |
| Brensholmen and Slettnes | Norway | 2018/19 | 3 |
| Rousay and Fair Isle | Scotland | 2017/18 | 6 |
| Rousay and Fair Isle | Scotland | 2018/19 | 6 |
| Rousay and Fair Isle | Scotland | 2019/20 | 2 |
| Kongsfjorden | Svalbard | 2009/10 | 13 |
| Kongsfjorden | Svalbard | 2010/11 | 13 |
| Kongsfjorden | Svalbard | 2011/12 | 17 |
| Kongsfjorden | Svalbard | 2012/13 | 12 |
| Kongsfjorden | Svalbard | 2013/14 | 4 |
| Kongsfjorden | Svalbard | 2014/15 | 6 |
| Kongsfjorden | Svalbard | 2015/16 | 6 |
| Kongsfjorden | Svalbard | 2016/17 | 13 |
| Kongsfjorden | Svalbard | 2017/18 | 7 |
| Kongsfjorden | Svalbard | 2018/19 | 7 |

**Table S2.** Number of deployments, of 179 in total, of each geolocator type on individual Arctic Skuas by population.

| Geolocator type | | Faroe Islands | Norway | Scotland | Svalbard | Total |
| --- | --- | --- | --- | --- | --- | --- |
| Migrate Technology | C250 | 23 | 47 | 0 | 17 | 87 |
|  | C65 | 0 | 1 | 12 | 1 | 11 |
| British Antarctic Survey | mk | 0 | 0 | 0 | 0 | 6 |
|  | mk13 | 0 | 0 | 0 | 13 | 15 |
|  | mk15 | 0 | 6 | 0 | 16 | 29 |
|  | mk18h | 0 | 0 | 0 | 0 | 1 |
|  | mk9 | 0 | 0 | 0 | 11 | 11 |
| Biotrack | mk3006 | 0 | 7 | 0 | 8 | 16 |
|  | mk4083 | 0 | 0 | 0 | 0 | 1 |

**Table S3.** Number of tracks per geolocator wet/dry recording mode and per population.

| Population | 3 seconds every  5 minutes | 3 seconds every  10 minutes | Every 6 seconds and  recorded on  change of state | No wet/dry  data collected | Total |
| --- | --- | --- | --- | --- | --- |
| Scotland | 0 | 10 | 0 | 4 | 14 |
| Faroe Islands | 0 | 15 | 12 | 0 | 27 |
| Norway | 4 | 23 | 73 | 2 | 102 |
| Svalbard | 0 | 74 | 0 | 24 | 98 |
| Total | 4 | 122 | 85 | 30 | 241 |

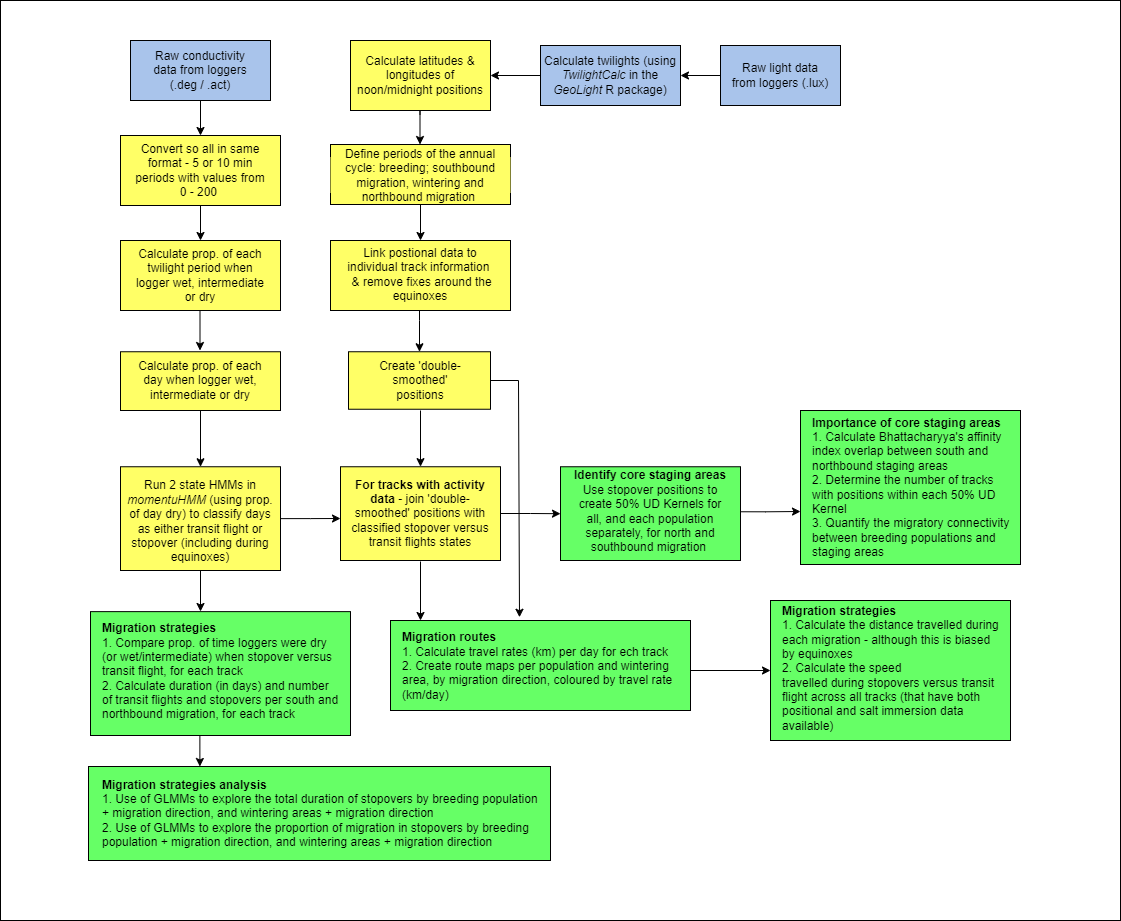

**Figure S1.** Flow chart showing how the data from the geolocators was processed and analysed. Blue boxes represent to input data, yellow boxes refer to processing steps and green boxes to analytical steps.

| Population | Tracks | Southbound | | Northbound | |
| --- | --- | --- | --- | --- | --- |
|  |  | Proportion of tracks | Mean proportion of migration | Proportion of tracks | Mean proportion of migration |
| Svalbard | 98 | 0.90 | 0.44 | 0.00 | 0.00 |
| Norway | 102 | 0.89 | 0.60 | 0.44 | 0.31 |
| Faroe Islands | 27 | 0.59 | 0.48 | 0.56 | 0.46 |
| Scotland | 14 | 0.79 | 0.51 | 0.71 | 0.36 |

**Table S4.** Proportion of tracks and mean proportion of migration duration, by breeding population, that overlapped with the equinoxes during south and northbound migration.

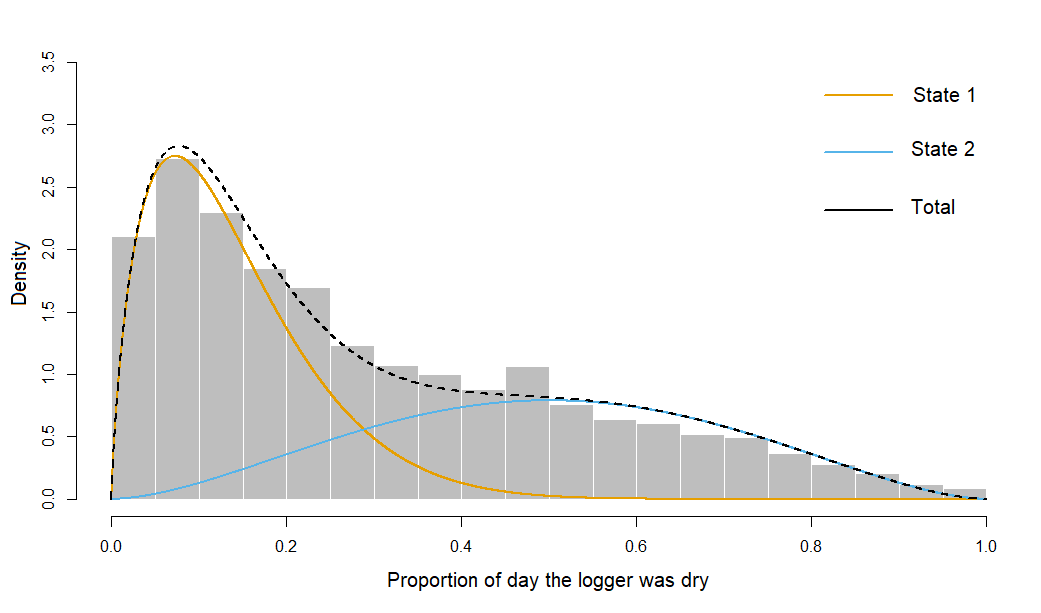

**Figure S2.** Classification output from the two-state hidden Markov model (HMM) with a single data stream, the proportion of each day that was dry. Positions classified as state 1 were considered at stopovers, associated with days where a lower proportion of the day was recorded as dry. Positions classified as state 2 were considered on transit flights, associated with days where a greater proportion of the day was dry.

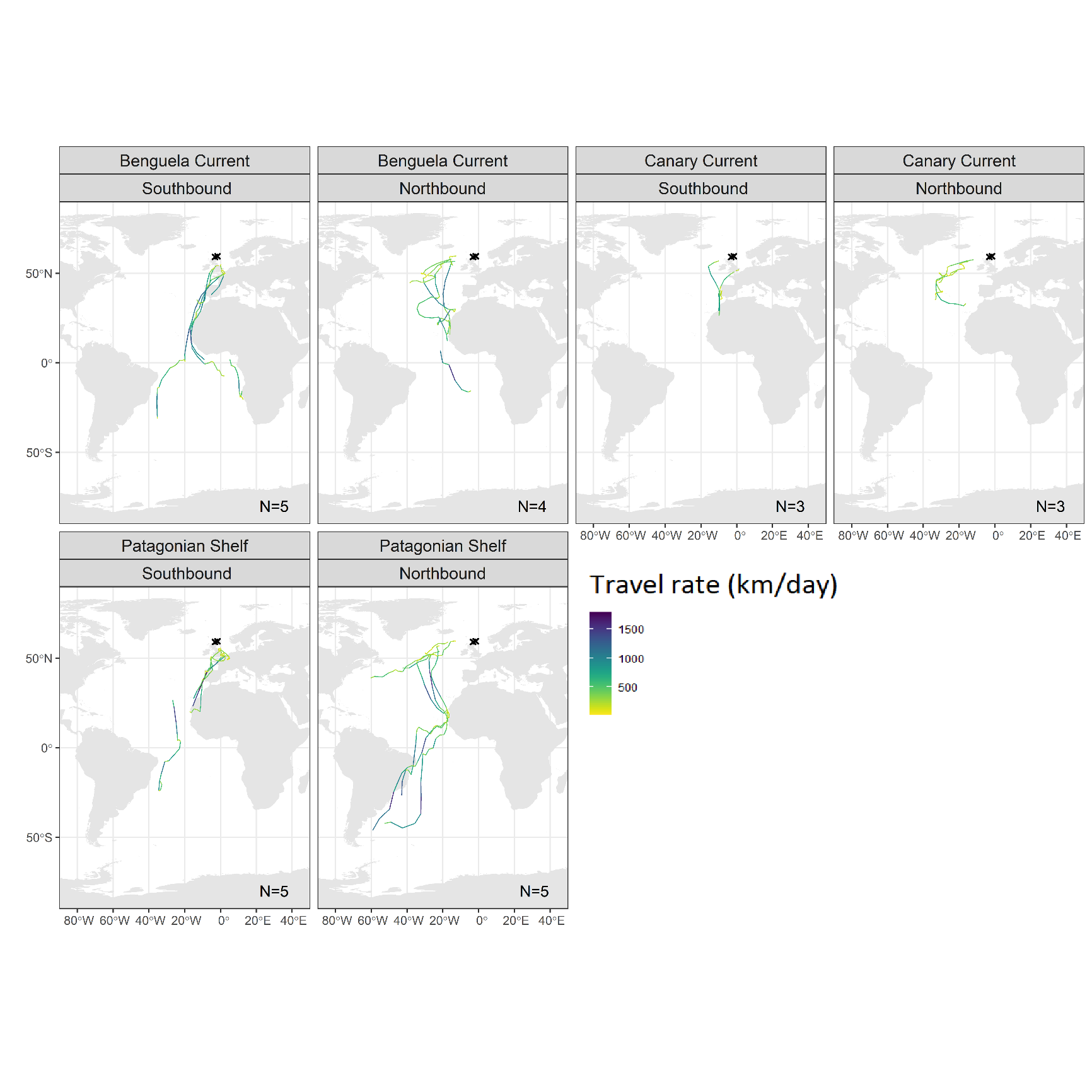

a) Scotland

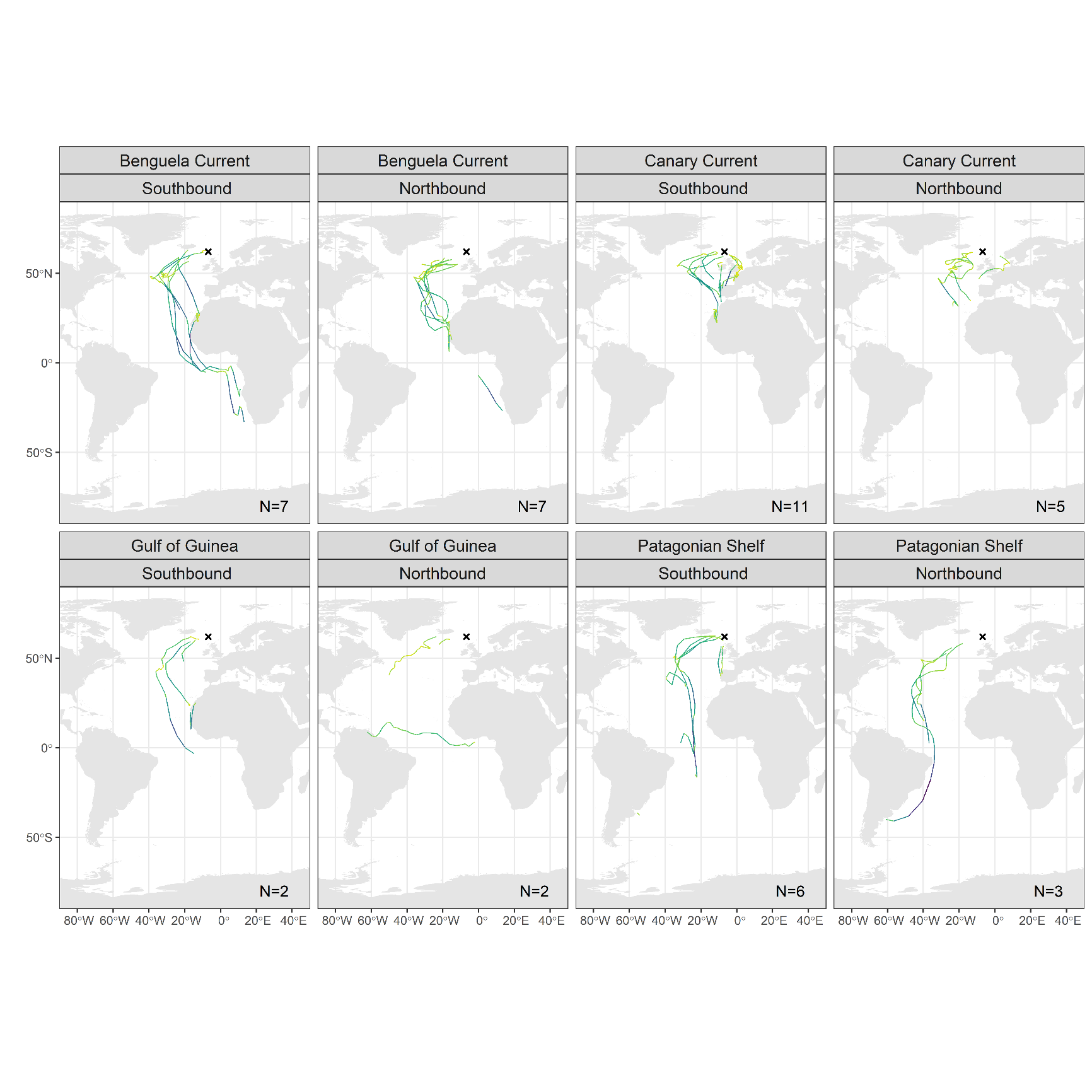

b) Faroe Islands

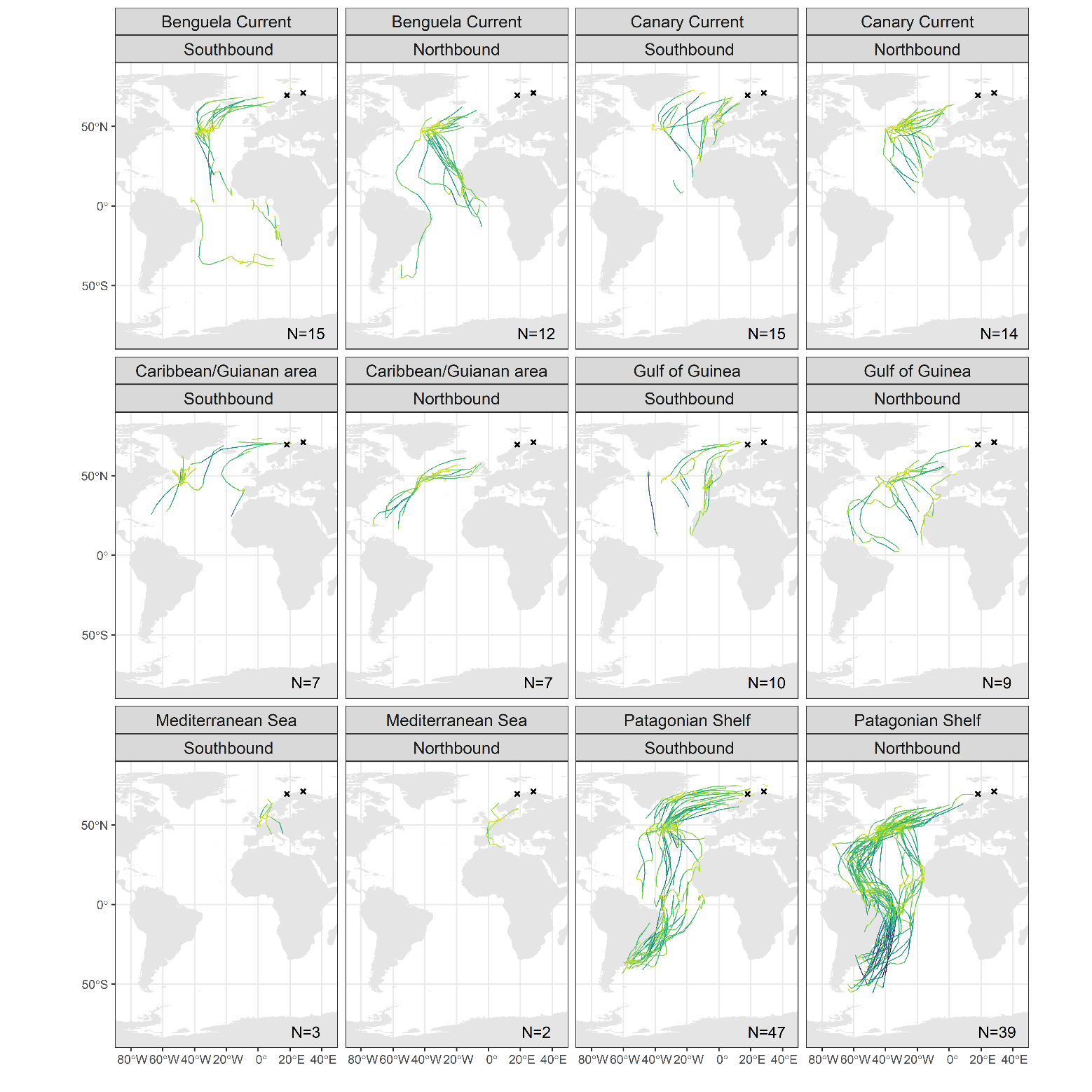

c) Norway

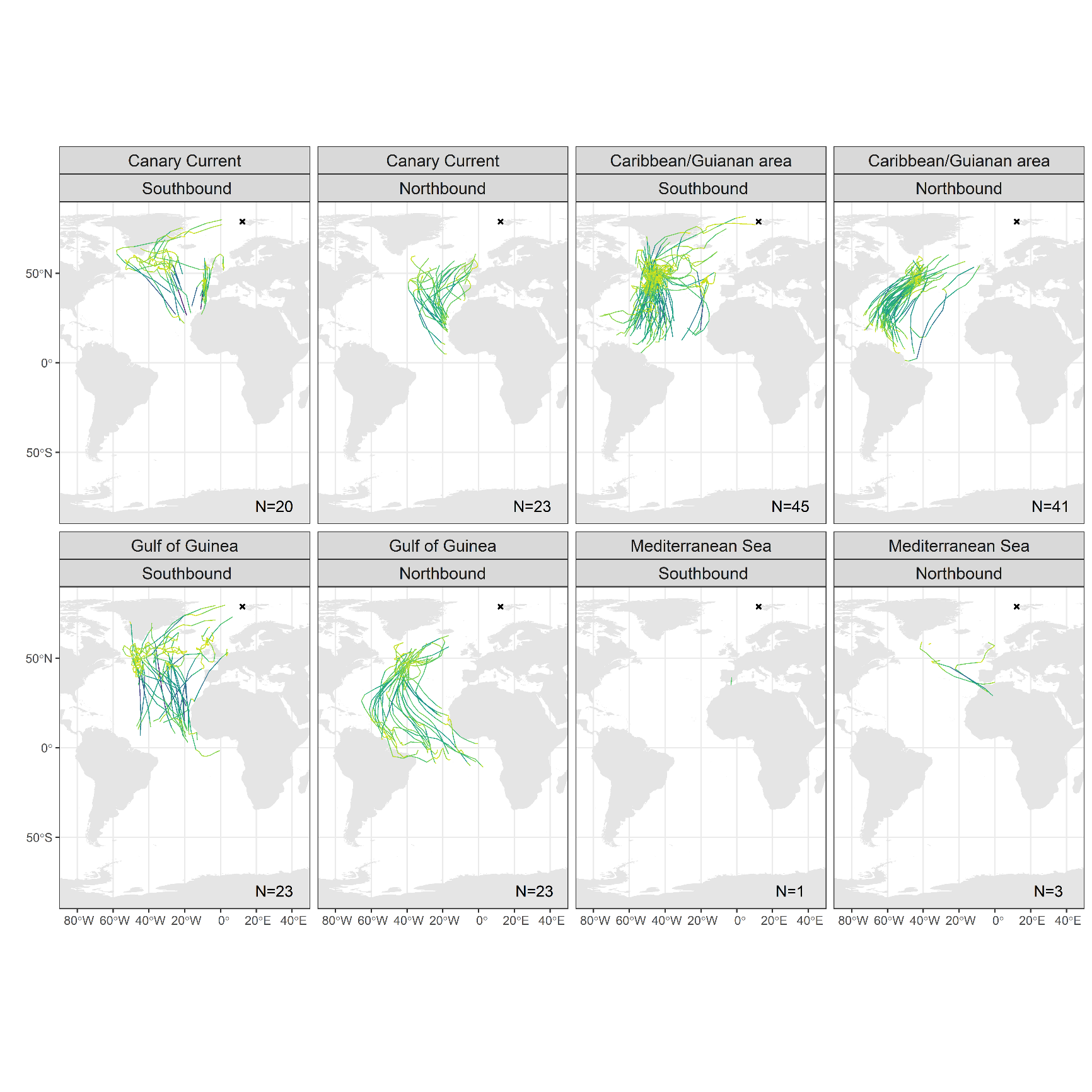
**Figure S3**. Smoothed migration routes of Arctic Skuas from a) Scotland, b) Faroe Islands, c) Norway and d) Svalbard during southbound and northbound migration, split by wintering area. The travel rate (km/day) of sections of tracks are shown to indicate areas where skuas were likely flying straight through (high rates of travel: dark green to purple) compared to those where individuals were foraging or resting on the water (low rates of travel: yellow to light green). Crosses depicts breeding colonies. N refers to number of tracks displayed for each migration period and wintering area. To visually identify areas that individuals migrated straight over (potentially indicating areas with lower productivity/foraging opportunities, Alerstam, 2009) compared to areas where individuals travelled more slowly and stopped to forage/rest, we calculated travel rates per day by measuring the daily distance travelled between double smoothed positions, using the *disthaversine* function in the *Geosphere* R package (Hijmans 2019).

d) Svalbard

### Among population and year consistency in core staging areas

As we were interested in identifying the core staging areas of Arctic Skuas across populations and years we combined the data from all tracks with activity data. To check these staging areas were representative of those used at the population level we also created utilisation distributions (UD) kernels for each population, for both the south and northbound migration (Figure S4). We calculated the overlapping space use of the 50% UD kernels between populations, and to the overall UD kernel, using Bhattacharyya's affinity index (BA, Bhattacharyya, 1943; Fieberg and Kochanny, 2005) and the *kerneloverlaphr* function in *adehabitatHR* (Calenge et al. 2015). For 50% UD kernels, the BA index provides a measure of similarity between two UD kernels between 0 and 0.5, with 0 indicating no overlap and 0.5 being identical (Fieberg & Kochanny 2005).

The extent of BA between the population specific 50% UD kernels and the overall species-level 50% UD kernels varied by season, with higher BA in north than southbound migration (southbound: 0.20 ±SD 0.17; northbound: 0.37 ± 0.07; Figure S4, Table S4). There were some differences in staging areas among populations. This was particularly evident for Scotland where the main southbound staging hotspot covered an area from the North Sea along the Iberian Peninsula, attributed to individuals travelling south along this route rather than via the mid-Atlantic ridge, which was the strategy used by most individuals from the more northerly populations of Faroe Islands, Norway and Svalbard. The main southbound staging area in the mid-Atlantic for Svalbard was also further east than that for the Faroe Islands.

Data on Arctic Skua migrations were obtained over multiple years, however we pooled the data across years to create the overall species-level UD kernel. To check whether the skuas were consistent in their core staging areas across years at the population level we created UD kernels for Svalbard and Norway for all years where at least 10 individuals were tracked. BA overlap between year-specific core 50% UD kernels were compared to the core 50% UD kernel for all years combined, for the Svalbard and Norway populations separately. For both Norway and Svalbard, there was high BA between the year-specific core UD kernels and the core UD kernel for each population of all years combined, across both migrations (Table S5, Figures S5 and S6).

**Figure S4**. Population-level core staging 50% utilisation distributions (UD) for a) southbound migration and b) northbound migration, based on positions classified as stopover locations, for Scotland (10 tracks), Faroe Islands (27 tracks), Norway (98 tracks) and Svalbard (74 tracks). Points indicate stopover locations (classified from 2 state HMM using saltwater immersion data, see main text for details) within and outwith the core 50% staging areas. Stars depict the breeding colonies.

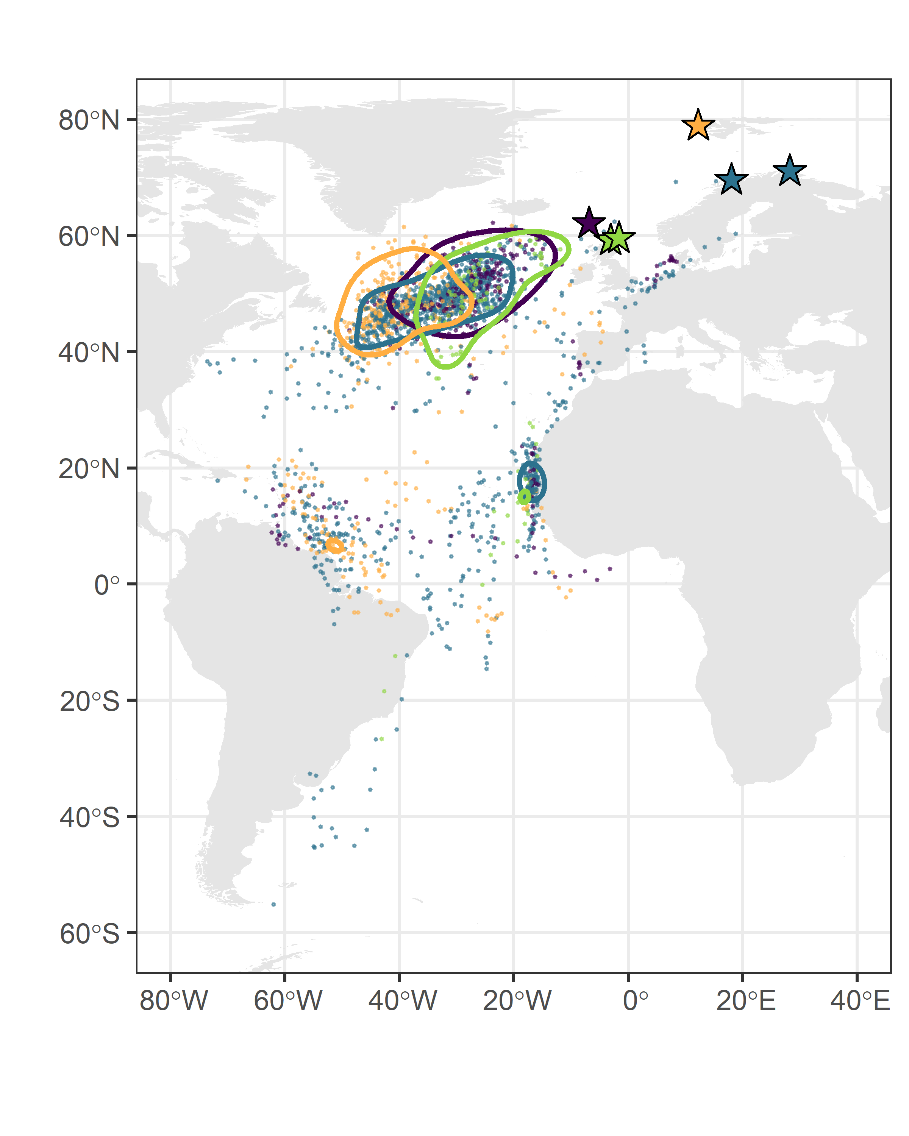

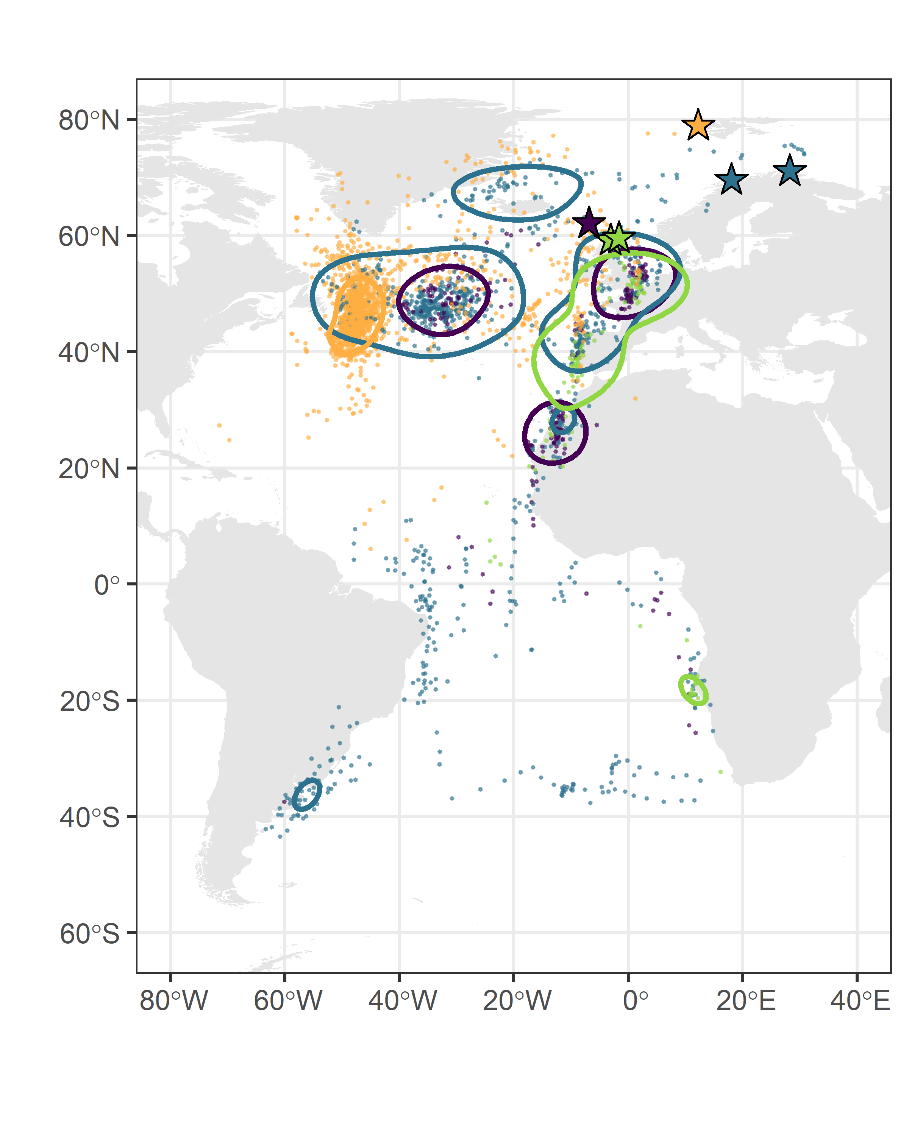

a)

b)

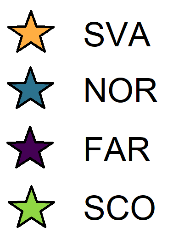

**Table S5.** Bhattacharyya's affinity (BA) in staging area 50% utilisation distributions, based on positions classified as stopover locations, compared among populations and to all tracks from the four populations combined, for south and northbound migration. Maximum BA value is 0.5.

|  | All tracks | Faroe Islands | Norway | Scotland | Svalbard |
| --- | --- | --- | --- | --- | --- |
| Southbound |  |  |  |  |  |
| All tracks |  | 0.12 | 0.29 | 0.01 | 0.38 |
| Faroe Islands |  |  | 0.26 | 0.16 | 0.00 |
| Norway |  |  |  | 0.18 | 0.13 |
| Scotland |  |  |  |  | 0.00 |
| Svalbard |  |  |  |  |  |
| Northbound |  |  |  |  |  |
| All tracks |  | 0.35 | 0.47 | 0.31 | 0.36 |
| Faroe Islands |  |  | 0.35 | 0.43 | 0.17 |
| Norway |  |  |  | 0.30 | 0.33 |
| Scotland |  |  |  |  | 0.12 |
| Svalbard |  |  |  |  |  |

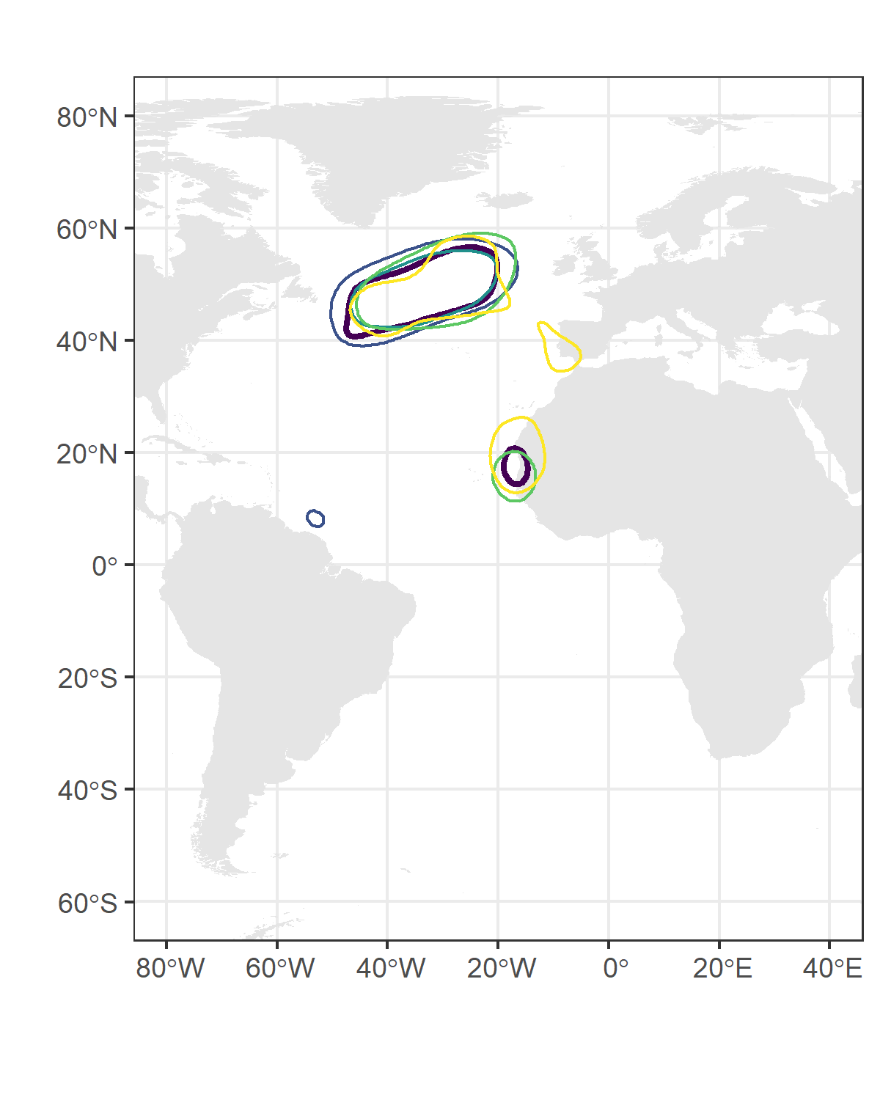

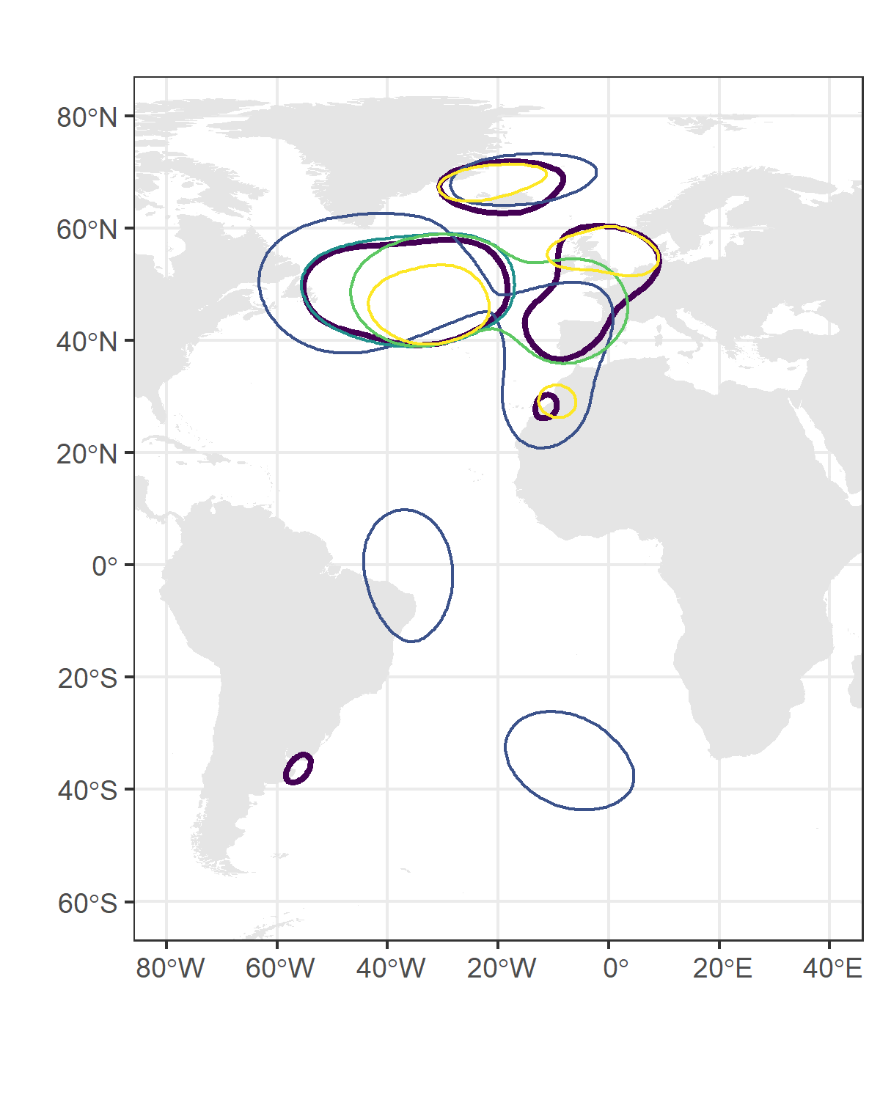

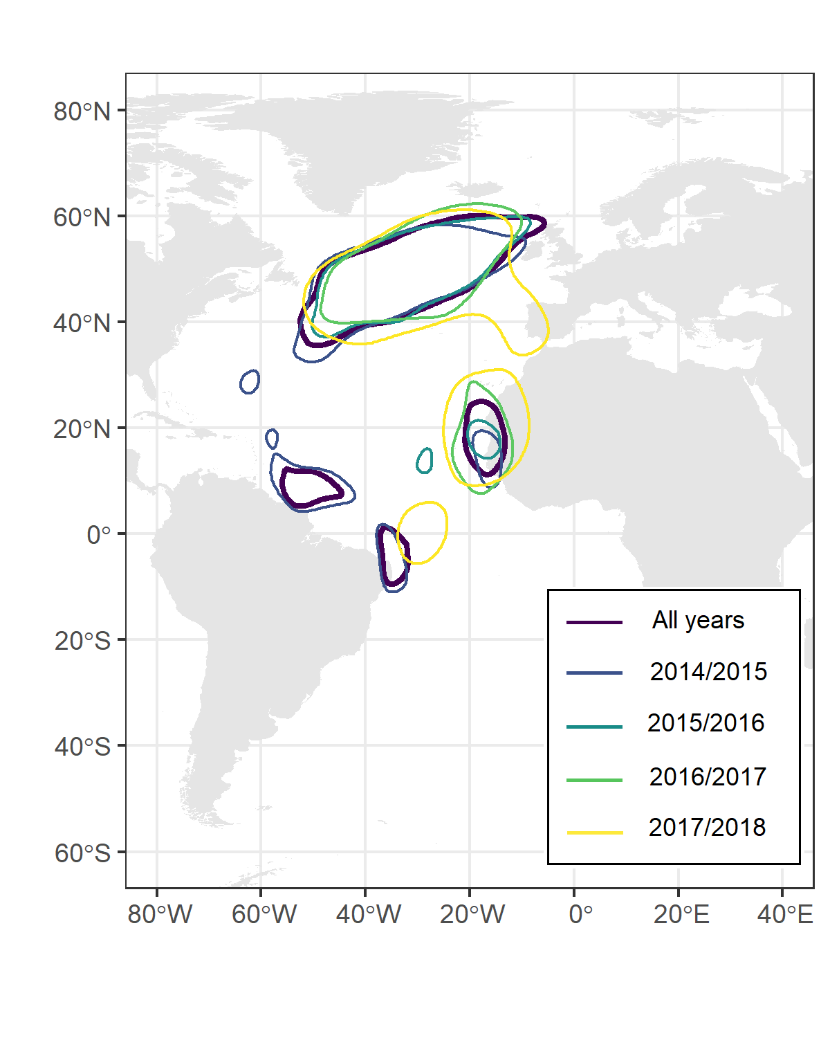

a)

b)

**Figure S5.** Norway core 50% utilisation distribution (UD) for all tracks and years (N =102), for a) south and b) northbound migration, based on positions classified as stopover locations, compared to the 50% core for individuals tracked during 2014/2015 (N = 26); 2015/2016 (N = 26); 2016/2017 (N = 16); and 2017/2018 (N = 12).

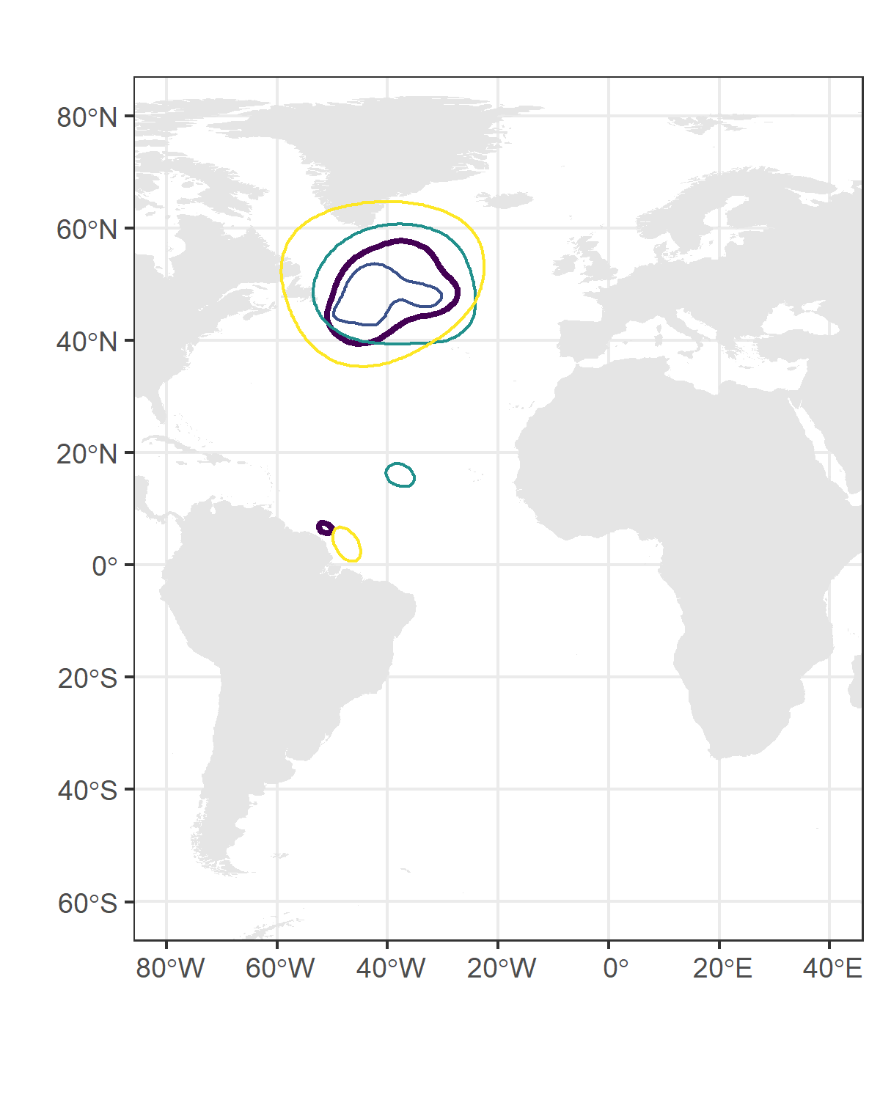

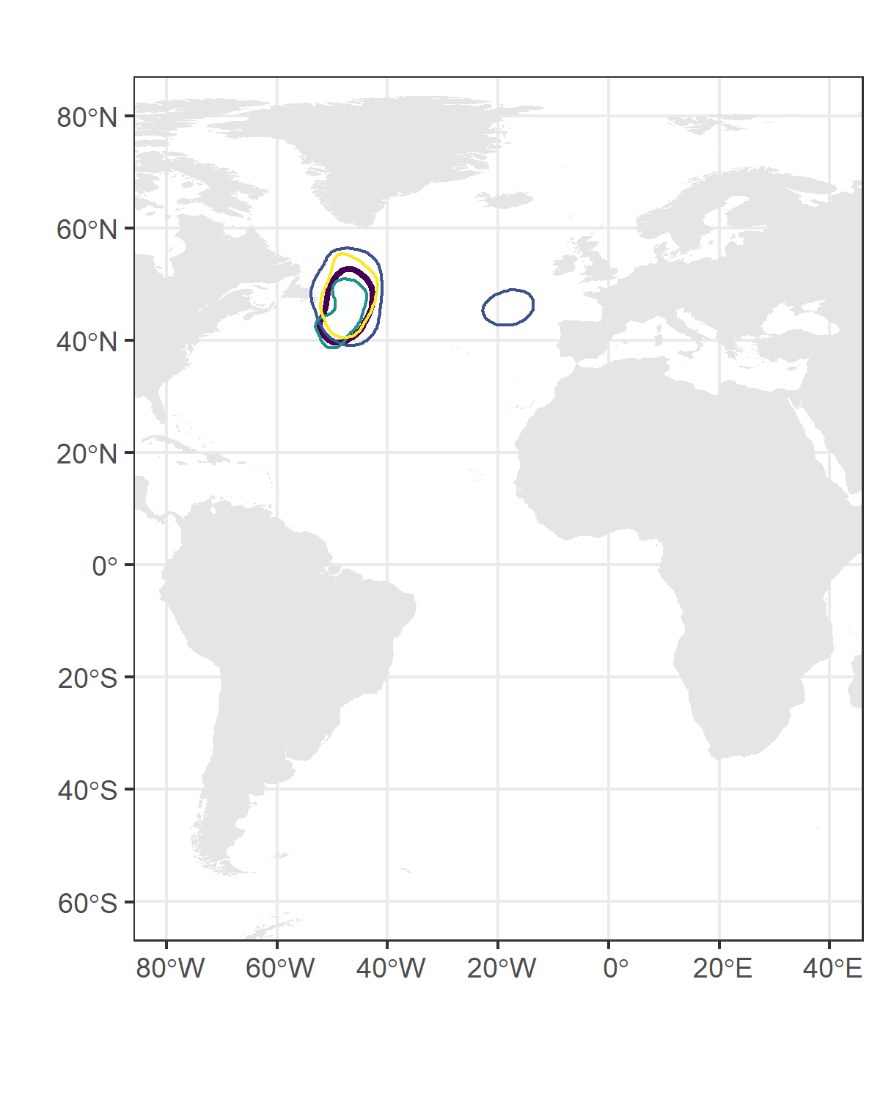

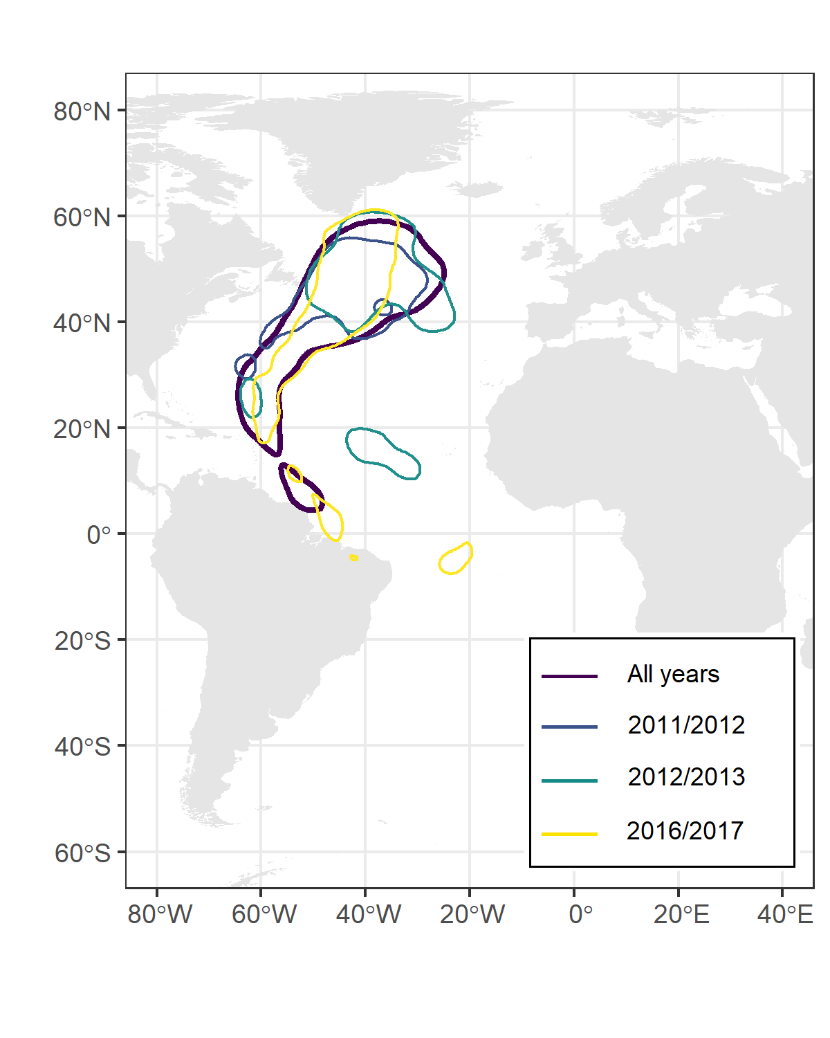

a)

b)

**Figure S6.** Svalbard core 50% utilisation distribution (UD) for all tracks and years (N =98), for a) south and b) northbound migration, based on positions classified as stopover locations, compared to the 50% core for individuals tracked during 2011/2012 (N = 17); 2012/ 2013 (N = 12); and 2016/2017 (N = 13).

**Table S6.** The extent of overlap, measured using Bhattacharyya's affinity (BA), between the population level 50% utilisation distribution (UD), based on positions classified as stopover locations, compared for all years combined and each year with more than 10 tracks was relatively high for both north and southbound migration, for a) Norway and b) Svalbard. Maximum BA value is 0.5.

| a) Norway | 2014/15 | 2015/16 | 2016/17 | 2017/18 |
| --- | --- | --- | --- | --- |
| No. of tracks | 26 | 26 | 16 | 12 |
| Southbound | 0.27 | 0.41 | 0.40 | 0.36 |
| Northbound | 0.42 | 0.46 | 0.41 | 0.37 |

| b) Svalbard | 2011/12 | 2012/13 | 2016/17 |
| --- | --- | --- | --- |
| No. of tracks | 17 | 12 | 13 |
| Southbound | 0.36 | 0.43 | 0.44 |
| Northbound | 0.38 | 0.40 | 0.34 |

**Table S7.** The number and proportion (in parenthesis) of individuals (which had tracks with saltwater immersion data and migratory timing details^1^) that had stopovers / any positions (stopovers and transit flights) within each core staging area (50% Kernel utilisation distributions) for all populations and both the south and northbound migration (Figure 1). The staging area codes match those in Figure 1 and 2. A proportion of individuals (southbound: 0.18; northbound: 0.19) were not recorded in the core staging areas likely due to missing data around the equinoxes, whilst some individuals visited more than one staging area during a migration period therefore the sum of individuals associated with staging areas differs to the total number of individuals.

| Staging area | Area (km^2^) | Centroid longitude | Centroid latitude | Faroe Islands | Norway | Scotland | Svalbard | Total |
| --- | --- | --- | --- | --- | --- | --- | --- | --- |
| Southbound |  |  |  |  |  |  |  |  |
| S1 | 3050000 | -41.41659 | 48.90871 | 7 / 12  (0.30 / 0.52) | 25 / 28  (0.48 / 0.54) | 0 / 0  (0.00 / 0.00) | 27 / 29  (0.79 / 0.85) | 59 / 69  (0.50 / 0.58) |
| S2 | 78400 | -8.528186 | 44.66028 | 3 / 4  (0.13 / 0.17 | 3/ 6  (0.06 / 0.12) | 1 / 1  (0.10 / 0.10) | 3 / 4  (0.09 / 0.12) | 10 / 15  (0.08 / 0.13) |
| S3 | 43700 | -18.15046 | 45.66223 | 0 / 1  (0.00 / 0.04) | 0 / 0  (0.00 / 0.00) | 0 / 0  (0.00 / 0.00) | 2 / 2  (0.06 / 0.06) | 2 / 3  (0.02 / 0.03) |
| S4 | 10200 | 1.305396 | 53.41426 | 2 / 2  (0.09 / 0.09) | 0 / 1  (0.00 / 0.02) | 1 / 1  (0.10 / 0.10) | 1 / 1  (0.03 / 0.03) | 4 / 5  (0.03 / 0.04) |
| None | NA | NA | NA | 11 / 5  (0.48 / 0.22) | 24 / 17  (0.46 / 0.33) | 8 / 8  (0.80 / 0.80) | 4 / 2  (0.12 / 0.06) | 47 / 32  (0.39 / 0.27) |
| Total individuals | - | - | - | 23 | 52 | 10 | 34 | 119 |
| Northbound |  |  |  |  |  |  |  |  |
| N1 | 2630000 | -33.88603 | 49.12961 | 13 / 13  (0.76 / 0.76) | 49 / 50  (0.94 / 0.96) | 4 / 4  (0.50 / 0.50) | 27 / 31  (0.77 / 0.89) | 93 / 98  (0.83 / 0.88) |
| N2 | 223000 | -16.83709 | 17.30656 | 3 / 3  (0.18 / 0.18) | 8 / 8  (0.15 / 0.15) | 4 / 4  (0.50 / 0.50) | 1 / 1  (0.003 / 0.03) | 16 / 16  (0.14 / 0.14) |
| None | NA | NA | NA | 4 / 4  (0.24 / 0.24) | 3/ 2  (0.06 / 0.04) | 0 / 0  (0.00 / 0.00) | 8 / 4  (0.23 / 0.11) | 15 / 10  (0.13 / 0.09) |
| Total individuals | - | - | - | 17 | 52 | 8 | 35 | 112 |

^1^ To identify the core staging areas using the 50% UD kernels we could only use fixes identified as stopovers outside the equinoxes where we also had positional data. Therefore, the use of the core staging areas, and number of individuals using each, is likely to be an underestimation as for most individuals the south and northbound migration coincided to some extent with the equinoxes. Given that the mid-Atlantic staging areas covered a large area of the Atlantic, we were unable to use the more reliable longitude of tracks to estimate which additional individuals used this staging area during the equinox.

**Table S8.** Estimated mean (± SD) number, duration and total duration of transit flights and stopovers taken by individual Arctic Skuas during, southbound and northbound migration split by breeding population, ordered from the highest to lowest latitude.

| Breeding area | Svalbard | | Norway | | Faroe Islands | | Scotland | |
| --- | --- | --- | --- | --- | --- | --- | --- | --- |
| Activity | Transit flights | Stopovers | Transit flights | Stopovers | Transit flights | Stopovers | Transit flights | Stopovers |
| Southbound migration | |  |  |  |  |  |  |  |
| Number | 2.73  (± 1.09) | 1.96  (± 1.04) | 2.80  (± 1.08) | 2.38  (± 1.14) | 2.37  (± 1.08) | 2.05  (± 0.86) | 2.50  (± 1.72) | 2.63  (± 1.6) |
| Duration | 5.04  (± 3.48) | 14.30  (± 16.3) | 7.78  (± 6.82) | 9.79  (± 10.4) | 7.19  (± 6.93) | 9.35  (± 10.1) | 8.00  (± 8.45) | 8.36  (± 6.76) |
| Total duration | 13.76  (± 4.50) | 27.93  (± 20.43) | 21.81  (± 10.97) | 23.27  (± 15.53) | 17.04  (± 10.54) | 19.14  (± 14.81) | 20.00  (± 13.58) | 22.00  (± 12.35) |
| Distance (km)^1^ | 8230 (± 1069) | | 11629 (± 3440) | | 8595 (± 3732) | | 9911 (± 3193) | |
| Northbound migration | | |  | |  | |  | |
| Number | 1.45  (± 0.65) | 1.32  (± 0.62) | 2.67  (± 0.92) | 2.01  (± 0.92) | 2.15  (± 0.88) | 1.85  (± 0.81) | 2.00  (± 0.87) | 1.88  (± 0.83) |
| Duration | 4.81  (± 2.83) | 7.27  (± 3.78) | 5.81  (± 4.65) | 9.62  (± 6.49) | 7.16  (± 6.09) | 10.90  (± 7.72) | 7.89  (± 7.04) | 9.27  (± 6.54) |
| Total duration | 6.97  (± 3.36) | 9.60  (± 5.11) | 15.49  (± 7.45) | 19.36  (± 9.08) | 15.40  (± 8.34) | 20.25  (± 12.87) | 15.78  (± 8.73) | 17.38  (± 7.91) |
| Distance (km)^1^ | 9083 (± 2690) | | 16117 (± 6059) | | 11478 (± 5980) | | 14269 (± 5793) | |

^1^ For each track, we calculated the great circle distance travelled during southbound and northbound migration from the breeding colony to the wintering area, including the direct line distance between the last and first location either side of the equinoxes, using the *disthaversine* function in the *Geosphere* R package (Hijmans 2019) on the double smoothed positions. Distances only provide a broad indication of the actual distances travelled by individuals given the error around raw geolocator positional fixes and due to the gaps around the equinoxes.

**Table S9.** Estimated mean (± SD) number, duration and total duration of transit flights and stopovers taken by individual Arctic Skuas during, southbound and northbound migration, split by wintering location, ordered from closest to furthest distance from the breeding populations.

| Wintering area | Mediterranean Sea | | Canary Current | | Caribbean region | | Gulf of Guinea | | Benguela Current | | Patagonian Shelf | |
| --- | --- | --- | --- | --- | --- | --- | --- | --- | --- | --- | --- | --- |
| Southbound migration | | | | | | | | | | | | |
| Activity | Migrant flights | Stopovers | Migrant flights | Stopovers | Migrant flights | Stopovers | Migrant flights | Stopovers | Migrant flights | Stopovers | Migrant flights | Stopovers |
| Number | 2.17  (± 0.75) | 1.80  (± 0.84) | 2.04  (± 0.88) | 1.63  (± 0.79) | 2.84  (± 1.25) | 1.93  (± 1.24) | 2.42  (± 0.76) | 1.80  (± 0.48) | 3.04  (± 1.27) | 2.63  (± 1.13) | 3.24  (± 1.00) | 2.87  (± 1.08) |
| Duration | 4.23  (± 3.09) | 8.67  (± 10.85) | 4.77  (± 3.56) | 9.94  (± 10.00) | 5.01  (± 3.29) | 19.20  (± 20.06) | 5.71  (± 4.50) | 14.13  (± 15.41) | 9.33  (± 8.14) | 8.40  (± 5.57) | 8.49  (± 7.34) | 7.59  (± 4.83) |
| Total  duration | 9.17  (± 6.18) | 15.60  (± 11.44) | 9.74  (± 5.43) | 16.21  (± 11.69) | 14.21  (± 4.21) | 37.00  (± 23.80) | 13.81  (± 6.68) | 25.43  (± 19.18) | 28.36  (± 7.19) | 22.04  (± 12.66) | 27.47  (± 7.37) | 21.78  (± 9.63) |
| Distance (km)^1^ | 4193 (± 561) | | 6432 (± 1417) | | 8432 (± 464) | | 9055 (± 766) | | 11296 (± 902) | | 14400 (± 851) | |
| Northbound migration | | | | | | | | | | | | |
| Number | 1.75  (± 0.50) | 1.75  (± 0.50) | 1.82  (± 0.80) | 1.37  (± 0.72) | 1.43  (± 0.61) | 1.29  (± 0.53) | 2.07  (± 0.87) | 1.63  (± 0.84) | 2.55  (± 0.74) | 2.14  (± 0.65) | 2.84  (± 1.07) | 2.28  (± 0.96) |
| Duration | 3.14  (± 1.68) | 7.86  (± 4.91) | 4.23  (± 2.93) | 10.95  (± 8.26) | 4.10  (± 2.40) | 8.75  (± 6.08) | 5.14  (± 3.23) | 11.14  (± 7.50) | 9.32  (± 8.14) | 8.40  (± 5.57) | 7.18  (± 6.02) | 7.78  (± 5.16) |
| Total  duration | 5.50  (± 1.73) | 13.75  (± 4.57) | 7.71  (± 3.87) | 14.97  (± 9.74) | 5.86  (± 2.82) | 11.29  (± 9.65) | 10.67  (± 5.28) | 18.15  (± 12.65) | 18.59  (± 5.32) | 20.38  (± 6.74) | 20.37  (± 5.76) | 17.72  (± 7.62) |
| Distance (km)^1^ | 6422 (± 1765) | | 7145 (± 2097) | | 8804 (± 1926) | | 12152 (± 2743) | | 15506 (± 3667) | | 20748 (± 3449) | |

^1^ For each track, we calculated the great circle distance travelled during southbound and northbound migration from the breeding colony to the wintering area, including the direct line distance between the last and first location either side of the equinoxes, using the *disthaversine* function in the *Geosphere* R package (Hijmans 2019) on the double smoothed positions. Distances only provide a broad indication of the actual distances travelled by individuals given the error around raw geolocator positional fixes and due to the gaps around the equinoxes.

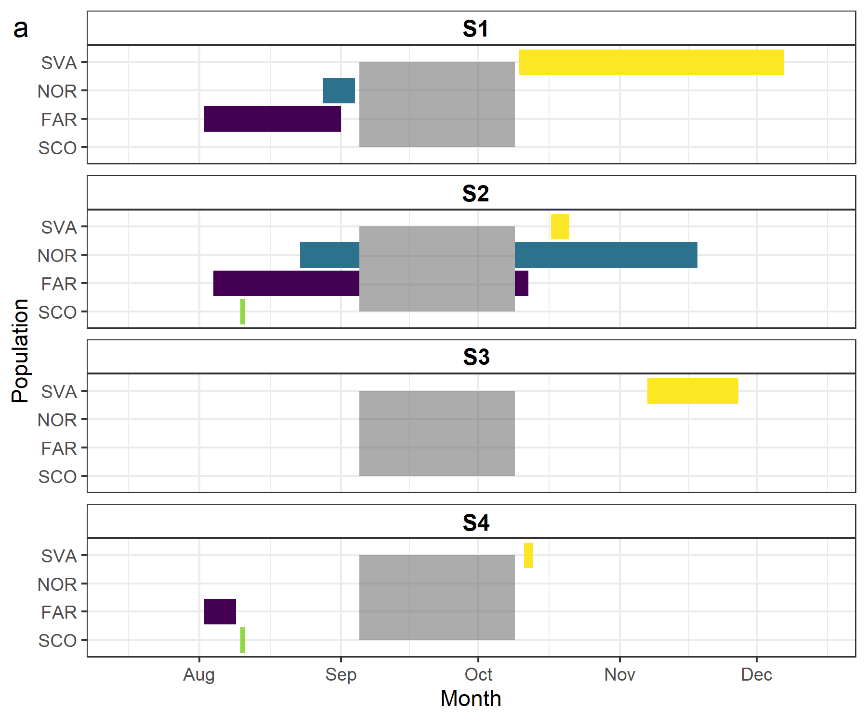

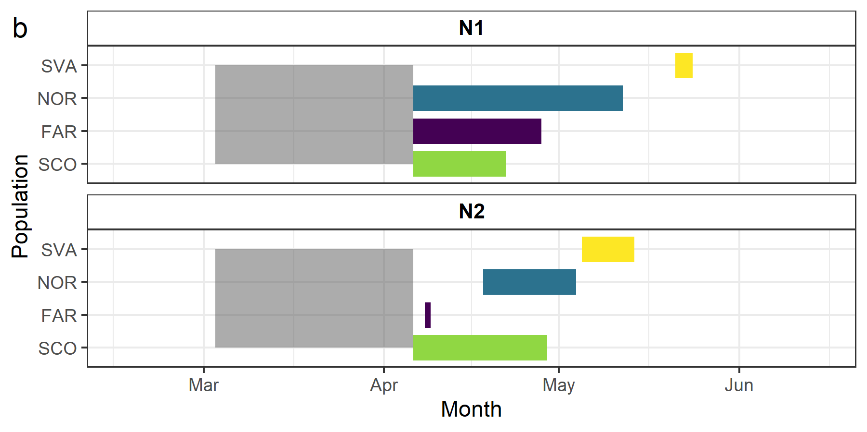

**Figure S7.** The first arrival date and last departure data of individuals to each staging area during a) southbound and b) northbound migration from each breeding population: Svalbard (yellow), Norway (blue), Faroe Islands (purple) and Scotland (green). The grey boxes cover the 17 days either side of the equinoxes (20 March and 22 September) where we lack positional data. Staging areas are labelled S1, S2 S3 and S4 for southbound migration and N1 and N2 for northbound migration (see Figure 1 and 2).

### **References**

Alerstam T (2009) Flight by night or day? Optimal daily timing of bird migration. J Theor Biol 258:530–536.

Bhattacharyya A (1943) On a measure of divergence between two statistical populations defined by their probability distributions. Bull Calcutta Math Soc 35:99–110.

Calenge AC, Dray S, Fortmann-roe S (2015) Package ‘adehabitat’.

Fieberg J, Kochanny CO (2005) Quantifying home-range overlap: the importance of the utilization distribution. J Wildl Manage 69:1346–1359.

Hijmans R (2019) Geosphere: Spherical Trigonometry. R package version 1.5-10.
